# Serum UCHL-1, GFAP, and NfL track tyrosine hydroxylase loss in substantia nigra in two Rat Models of Parkinson’s Disease

**DOI:** 10.64898/2026.07.14.738586

**Authors:** Isabel Soto, Robert McManus, Walter Navarrete, Isha Mhatre-Winters, Elle V. Rogers, Janiece Vancil, Kirby Doshier, Jason R. Richardson, Vicki A. Nejtek, Michael F. Salvatore

## Abstract

In Parkinson’s disease (PD), blood-based (BB) biomarkers ubiquitin c-terminal hydrolase L 1 (UCHL-1), glial fibrillary acidic protein (GFAP), and neurofilament light (NfL) correlate with motor or cognitive impairment. However, it is unclear if blood levels of these biomarkers represent changes in nigrostriatal neuron viability or dopamine (DA) signaling. In 6-OHDA and Pink1 knockout (KO) rat models that showed progressive loss of DA tissue and tyrosine hydroxylase (TH) protein, we quantified UCHL-1, GFAP, and NfL expression in striatum and substantia nigra (SN) at 7- and 28-days in the 6-OHDA model and 7- and 18-month old in Pink 1 KO. Substantial changes in all biomarkers occurred with TH loss in SN, but not striatum, in both models. UCHL-1 levels increased against remaining TH protein. Accordingly, serum UCHL-1 levels increased 25% at 28 days post-6-OHDA and 18-month old Pink1 KO. GFAP and NfL levels increased in SN 28 days post-6-OHDA and 18 month-old Pink1 KO. Serum GFAP levels increased 28 days post-6-OHDA and 18 month-old Pink1 KO. Serum levels of NfL increased 28 days post-6-OHDA, and in 18 month-old Pink1 KO and wild-type, without influence by genotype. Expression levels of each biomarker were greater in the SN vs striatum, suggesting the SN contributes greater quantities of biomarkers to the blood and reflect TH loss therein. Taken together, our preclinical results show alignment between serum levels of UCHL-1, GFAP, and NfL and loss of TH and DA in the SN. As such, these biomarkers may be relevant peripheral indicators of deficient nigrostriatal DA signaling, and reflect nigrostriatal function in PD.

## INTRODUCTION

The search for a dependable biomarker profile to positively identify Parkinson’s disease (PD) pathology and its severity has been ongoing for the past few decades. The α-synuclein aggregate or phosphorylated form has shown specificity or sensitivity to predict the presence of disease, but may not gauge disease severity (Pons et al., 2022; Pedersen et al., 2025). Brain-specific markers such as Ubiquitin C-terminal hydrolase L1 (UCHL-1), glial fibrillary acidic protein (GFAP), and neurofilament light chain (NfL) have repeatedly shown a relationship with neurodegeneration, including PD (Yamashita, 2023; Youssef, 2023). We have recently reported that serum levels of GFAP and NfL increase 4 weeks after 6-OHDA lesion induction. This increase is associated with major (>90%) loss of tyrosine hydroxylase (TH) and dopamine (DA) in the striatum, TH loss of ∼80% in the substantia nigra (SN), and neuronal loss of ∼80% (Kasanga et al., 2024). While these biomarkers have some utility to gauge symptom severity in PD (Su et al., 2012; Mollenhauer et al., 2020; Ye et al., 2021; Ygland Rödström et al., 2022; Lin et al., 2023; Tang et al., 2023; Youssef et al., 2023), it has yet to be established whether changes in these blood-based (BB) biomarkers correspond to changes in DA signaling or neuron viability. Thus, identifying if equivalent changes biomarker levels between the blood and the nigrostriatal pathway exist would answer to a long-standing knowledge gap as to whether BB biomarker levels are peripheral indicators of nigrostriatal degeneration or DA signaling.

UCHL-1 is both a deubiquitinating enzyme and a ubiquitin ligase, and highly prevalent in neurons (Cartier, 2012; Mi, 2021). The exact connection between UCHL-1 function and PD remains unknown, but it is possible that it may elevate free ubiquitin levels to promote clearance of protein aggregates such as alpha-synuclein (Kumar, 2017; Cartier, 2012; Ng, 2020). Conceivably, such an increase could act as a compensatory mechanism to clear increased protein aggregation, as occurs in PD progression. GFAP is a cytoskeletal protein of astrocytes to maintain morphology. Elevated GFAP is indicative of astrocyte activation in rodent models of PD (Kasanga et al., 2023a; Jeong et al., 2026). Expression of GFAP also increases with age, and increases in the blood and cerebrospinal fluid (CSF) of individuals with PD (Lin, 2023; Tang et al., 2023). Lastly, NfL is a component of neurofilaments found in the cytoskeleton of neurons and essential for maintaining neuronal shape and the structural integrity of the axon. Accordingly, blood levels of NfL correlate with both non-motor and motor deficits of PD (Ge et al., 2018; Backstrom et al., 2020; Benatar et al., 2020; Ma et al, 2021; Huang et al., 2022; Liu et al, 2022; Welton et al., 2022). While alterations in UCHL-1, GFAP, and NfL may not directly contribute to PD pathology, mounting evidence suggests they are viable markers of the severity of nigrostriatal degeneration and loss of DA signaling.

Longitudinal evaluation of these biomarkers in established preclinical PD models could clarify whether blood levels can estimate the extent of the loss of DA signaling in nigrostriatal pathway. Specifically, we determined how each biomarker may change in the striatum and substantia nigra (SN) against loss of DA and TH, and whether such change corresponded to a change in blood levels. To circumvent limitations of using only one PD model (Jackson-Lewis, 2012; Jagmag et al., 2016; Creed, 2018; Zhang, 2022), we used 2 rat PD models in longitudinal evaluations, which also increased rigor and translatability. Our 6-OHDA rat PD model produces rapid TH loss in striatum, but also a more protracted rate of TH+ cell loss in the SN at the same time (Kasanga et al., 2023a). This emulates the disparity of DA and TH loss in PD, being greater in striatum than in the SN upon diagnosis at onset of motor impairment (Bezard et al., 2001; Kordower et al., 2013). The genetic Pink1 knockout (KO) rat potentially represents a closer parallel to the slow rate of disease progression of human PD (Dave, 2014; Villeneuve, 2016; Cullen, 2018; Kelm-Nelson, 2021). Critically in both models, we reported that decreased locomotor activity was associated with loss of DA and TH in the SN, given lesion severity in the 6-OHDA model (Kasanga et al., 2023) or as a function of aging in the Pink1 KO model (Soto et al., 2024a). Thus, these two models provided comparable hypokinesia severity, but at a differential rate, that coincided with loss of DA markers in the SN, providing the opportunity to determine whether BB biomarkers consistently reflect nigrostriatal degeneration across, despite differing triggers of nigrostriatal decline different models of disease.

We recently reported 6-OHDA lesion increased GFAP and NfL serum levels (Kasanga et al., 2024), thus giving model-equivalent results for comparison to expression in CNS to determine levels of UCHL1, GFAP, and NfL in striatum and SN. In the Pink1 KO, we were able to make these blood to brain comparisons within the same rats to evaluate biomarker changes associated with progressive aging-related dopaminergic dysfunction. Our findings revealed that changes in expression profiles of these biomarkers consistently occurred in the SN, but not the striatum, between both models, increasing the predictive validity that these serum biomarkers, together, are an index of the severity of the loss of nigrostriatal DA signaling.

## METHODS

### Animals

Two different rat models of PD were used to provide insight into changes in biomarkers against the progression of loss of TH protein and DA in the striatum and SN; the unilateral 6-OHDA medial forebrain bundle lesion model (Kasanga et al., 2023; 2024) and the Pink1 KO (Soto et al., 2024a). Notably, in the 6-OHDA model, differences in the magnitudes of loss of DA and TH protein were observed in the striatal and SN tissue collected that resembled the dichotomy of loss exhibited in human PD (Kasanga et al, 2023), with greater loss in striatum than that in the SN (Kordower et al., 2013). Specifically, there was progressive loss of TH+ neurons, and in the SN, also progressive loss of tyrosine hydroxylase (TH) protein, and dopamine (DA) that occurred at a less rapid rate than in striatum (Kasanga et al., 2023) (Fig. 1). In the Pink1 KO, the older cohort of 18 months old had significantly less DA and TH compared to the younger cohort of 7 months old, but notably this loss was specifically in the SN and not in striatum (Fig. 2) (Soto et al., 2024).

**Figure 1.**
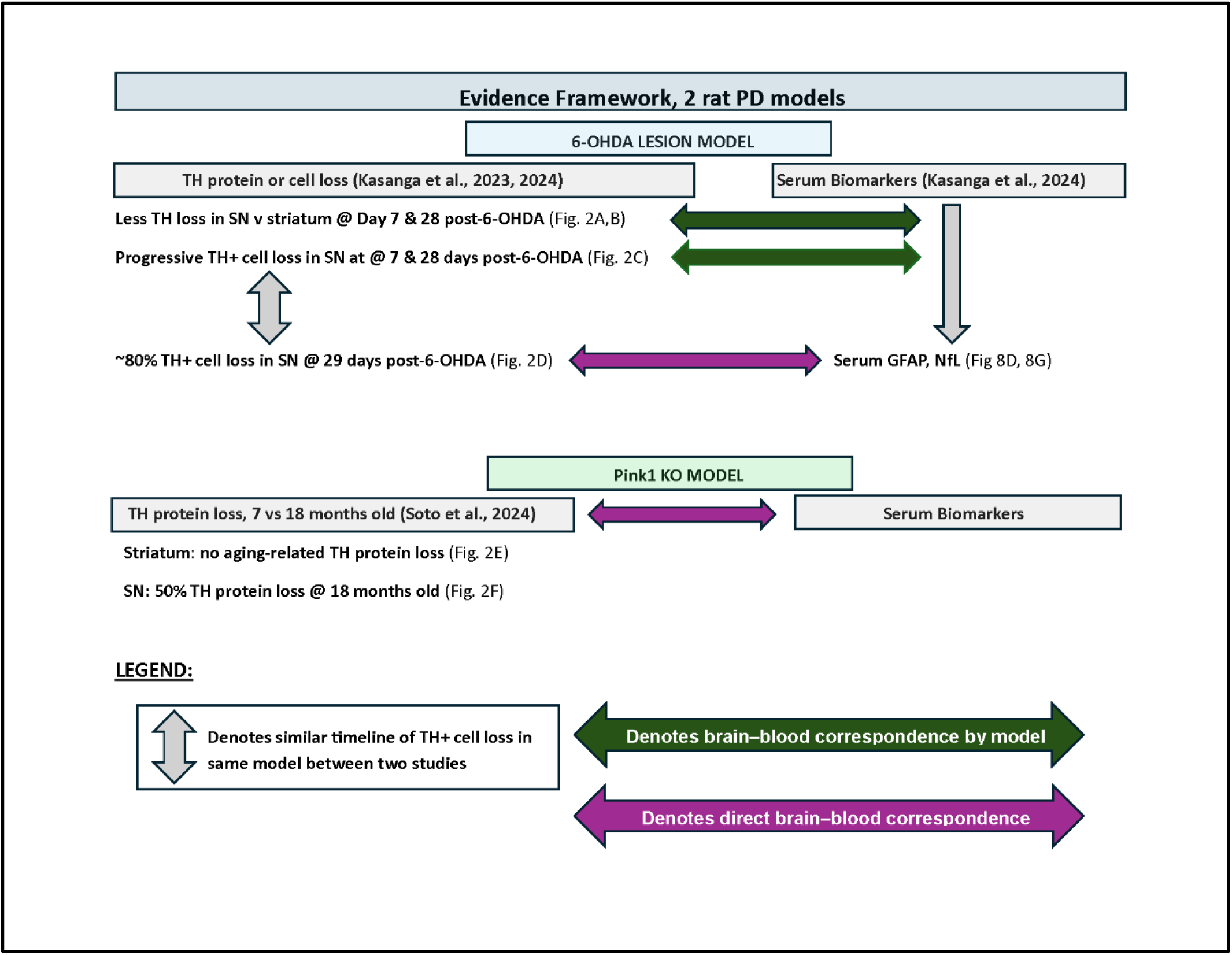
CNS tissue and blood source comparison from 3 studies. Flow chart representation of CNS tissue and serum sample relationships between the two rat PD models. The 6-OHDA model used produces gradual TH protein loss and TH+ cell loss in the SN over the course of 28 days. Serum was collected from one of these studies (Kasanga et al., 2024) and TH+ cell loss from this study was nearly identical to the study from which the CNS tissues were analyzed for biomarker expression profile and TH protein and DA loss were previously reported (Kasanga et al., 2023). In the Pink1 KO study, TH protein and DA loss were also progressive as a function of aging and reported (Soto et al., 2024). Blood was processed from this study and biomarkers in the CNS tissues were analyzed from remaining tissue samples. Thus, serum and CNS results are from the same study.

**Figure 2.**
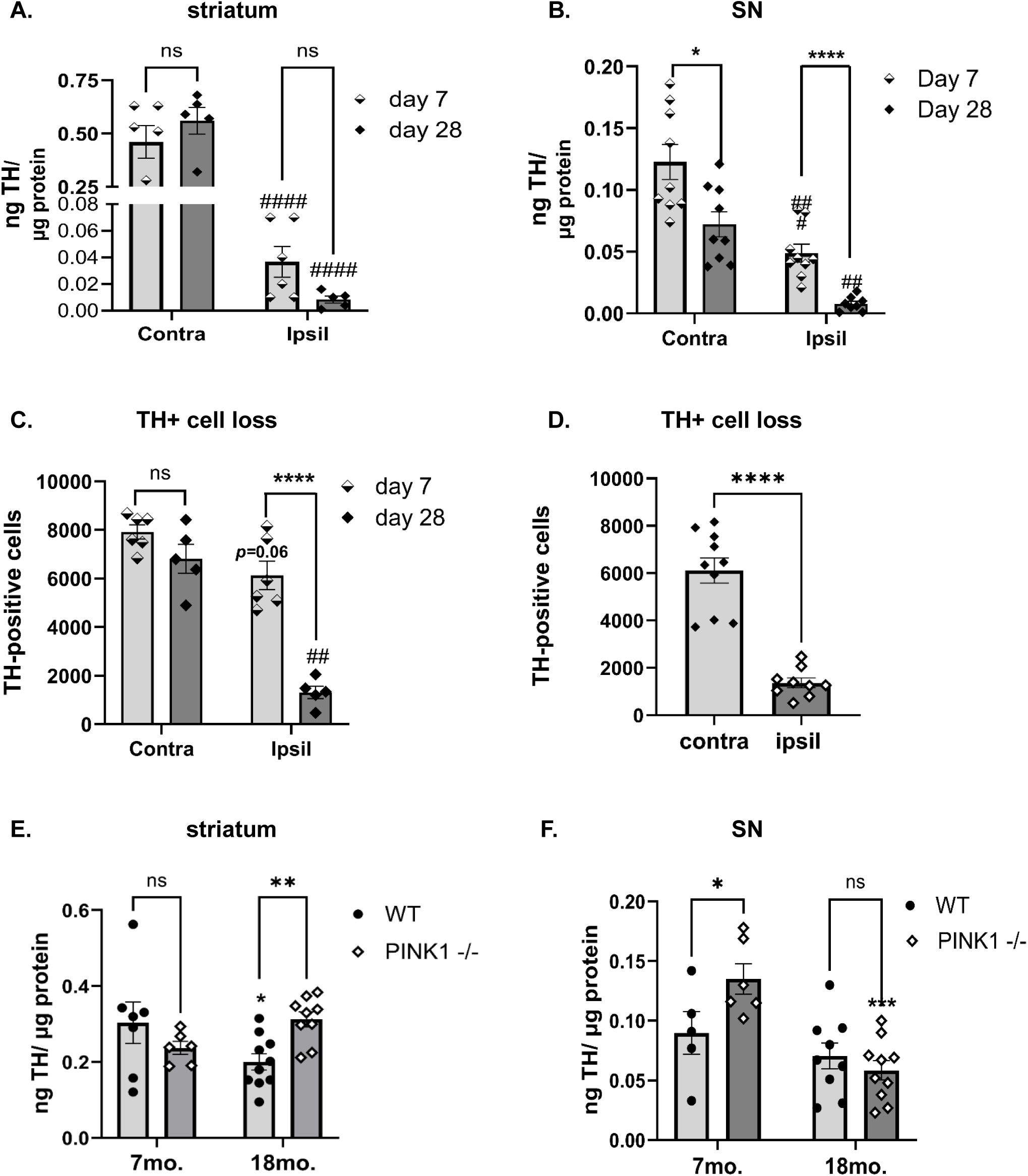
Progressive loss of nigrostriatal neurons vs loss of striatal tyrosine hydroxylase (TH) protein in the 6-OHDA model. Our 6-OHDA model emulates the differential loss of TH in striatum vs neuronal loss seen in human PD. At the time points evaluated in this study (7 and 28 days after induction of 6-OHDA lesion), loss of neurons in the SN occurred at a rate slower than TH protein loss in striatum. **A. striatal TH protein loss, 6-OHDA.** Loss of TH protein in striatum exceeded 90% by 7 days (after 6-OHDA lesion induction and was maximal with no further decrease by day 28. Contralateral to lesion (Contra) vs Ipsilateral to lesion (Ipsil). Day 7 (t=6.65, ^####^*p*<0.0001), Day 28 (t=7.91, ^####^*p*<0.0001). These results were similar to TH loss previously reported in striatum with this model (Kasanga et al., 2023), emulating the fidelity of this model to reproduce consistent TH protein loss. **B. nigral TH protein loss, 6-OHDA.** This previously reported result (Kasanga et al., 2023), shows progressive TH protein loss in the SN with this model 60% by day 7 and decreases further to ∼85% by day 28. Day 7 vs day 28, ipsil lesion (t=5.24, \*\*\*\**p<*0.0001), contra lesion (t=2.90, \**p*<0.011). **C. Progressive TH+ cell loss.** This previously reported results shows progression of TH+ cell loss increases as days after 6-OHDA lesion increased, with modest (∼20%) loss at day 7 (t=2.61, *p=* 0.057) vs ∼75% by day 28 (t=7.3, \*\*\*\**p*<0.0001). Lesion X days post-lesion (F(1,9)=13.3, *p*=0.005). Striatal TH loss results (presented above in panel A) are from same rats with TH+ loss shown in this figure. **D. TH+ cell loss 4 weeks after 6-OHDA.** This new set of results shows ∼80% TH+ cell loss by day 29 following 6-OHDA ((t=9.55, \*\*\*\**p<*0.0001), similar to that previously reported in Kasanga et al., 2023. Striatal TH loss was similar from this study (Kasanga et al., 2024), as compared to that reported in (Kasanga et al., 2023) and in panel A. **E. striatal TH protein loss, Pink1 KO.** TH protein levels decreased ∼30% only in the WT as a function of aging; 7 mo vs 18 mo (t=2.46, \**p*=0.020) whereas in the aged cohort alone, TH protein levels were significantly greater in the Pink1 KO vs WT (t=2.85 , \*\**p*=0.008)**. F. nigral TH protein loss, Pink1 KO.** TH protein levels in the WT were similar between age groups, unlike that in the striatum; 7 mo vs 18 mo (t=1.10, *p*=0.28). However, in the Pink1 KO, there was highly significant loss of TH at 18 mo vs 7 mo (t=4.71, *p*<0.0001). In the young cohort, TH protein levels were significantly greater in the Pink1 KO vs WT (t=2.37, \**p*=0.025).

In the 6-OHDA study, 65 male Sprague-Dawley rats, 3 months old, previously characterized for nigrostriatal neuron DA markers, were used to evaluate the biomarker profile in the SN and striatum in this timeline study, wherein DA tissue content and TH protein levels were previously determined at either 7 or 28 days after induction of 6-OHDA lesion. A sham-operation group served as a control group to evaluate whether differences in striatal or nigral tissue contralateral to lesion occurred as compared to any possible influence of the operation itself. In the Pink1 KO study, the groups consisted of young (7-month-old) WT (n=12; (Charles River) and Pink1 KO (n= 14; Envigo) male rats, and aged (18-month-old) WT (n=5 from Envigo and n=5 from Charles River) and Pink1 KO (n=10; Envigo) male rats. All rats were single-housed on a 12-hr reverse light/dark cycle with food and water provided ad libitum. For one month prior to the experimental procedures, rats were handled at least three times a week and acclimated to their environment.

Serum GFAP and NfL data were obtained from a previously published cohort consisting of non-exercise control group using 6-OHDA to induce nigrostriatal lesion (Kasanga et al., 2024). Thus, although the serum and CNS biomarker measurements were not collected from the same animals, both cohorts utilized the identical 6-OHDA model, experimental design, inclusion criteria, and time points, with comparable TH protein and neuronal loss between studies by 4 weeks after lesion induction, as illustrated in Figure 1. This allowed direct comparison of serum biomarker profiles with regional CNS biomarker expression while maintaining consistency in disease severity across cohorts. All experiments were performed under approved protocol by the institutional Animal Care and Use Committee (IACUC) at the University of North Texas Health Science Center and the Animal Care and Use Review Office (ACURO), Department of the Army, U.S. Army Medical Research and Development Council.

### 6-OHDA lesion induction

Rats were deeply anesthetized with 2- 3% isoflurane for stereotaxic surgery. Bregma was identified, and the medial forebrain bundle (mfb) was targeted in both hemispheres. Freshly-prepared 6-OHDA (16 µg in 4 µg/µl) in 0.02% ascorbic acid in sterile saline was infused into the mfb at coordinates relative to Bregma (in mm) AP -1.8, ML 2.0, DV -8.6 at a rate of 1 µl/min with a 26 gauge needle (Plastics One, Roanoke, VA). The needle was left in place for 10 min after infusion prior to withdrawal. The mfb on the side contralateral to lesion received an infusion of the same volume of 0.02% ascorbic acid in sterile saline (without 6-OHDA) to control for impact that infusion or tissue damage from the needle may have on expression of the proteins measured. A sham-operated group was included in the study, using identical time-points post-surgery. The sham-operated group received 0.02% ascorbic acid in sterile saline delivered into the mfb in one hemisphere. The mfb contralateral to the sham-operated side remained intact. After 6-OHDA lesion induction or sham-operation, the rats were randomly subdivided into two separate cohorts consisting of a 7-day post-lesion and 28-day post-lesion (or post sham-operation) group. This study design allowed us to determine if differences in UCHL-1, GFAP, or NfL expression were associated with the severity of nigrostriatal neuron loss, and if changes found contralateral to the lesion were due to loss of the neurons ipsilateral to lesion, and not due to sham-operation.

### Locomotor assessments

We previously reported locomotor results from both cohorts in this study (Kasanga et al., 2023; Soto et al., 2024a). For this study, these assessments provided the results needed to conduct statistical analyses on expression levels of the biomarkers on only the test subjects in the 6-OHDA cohort that exhibited a locomotor deficit; the criterion are >30% loss of forelimb use at 7 days after induction of nigrostriatal lesion, which strongly predicts eventual hypokinesia by 21 days after lesion induction (Kasanga et al., 2023).

### Quantification of TH protein and biomarkers in CNS

A critical component in the assessment of the biomarkers includes normalization against remaining TH protein, which is a gauge of biomarker expression against the progression of the impairment or loss of nigrostriatal neuron function in both models. The progressive loss of TH in the SN of the 6-OHDA model and the aging-related loss of TH in the SN of the Pink1 KO model have been previously reported in the tissues used in this biomarker assessment (Kasanga et al., 2023; Soto et al., 2024a). We quantified TH protein levels against calibrated TH standards, so that biomarker levels were normalized against TH protein (ng TH per µg total protein) remaining in the two models (Millipore, rabbit polyclonal, cat #AB152, 1:1000 dil).

For analysis of the biomarker profiles in CNS, we first optimized total protein load for each biomarker running a tissue standard curve at 5, 10, 20, and 30 µg of total protein. Nominal total protein loads were determined to be at 5 µg for UCHL-1 and GFAP, and 20 µg for NfL in both rat strains (Sprague-Dawley for 6-OHDA and Long-Evans for the Pink1 KO). The following sources of primary antibody were as follows; UCHL-1 (rabbit monoclonal, Cell Signaling cat#13179, 1:1000 dilution), GFAP (mouse monoclonal, Cell Signaling, cat #: 3670, 1:1000 dilution), and NfL (mouse monoclonal, Cell Signaling, cat #2835, 1:1000 dilution).

### Analysis of blood-based biomarkers

As previously reported, serum levels of NfL and GFAP were quantified from trunk blood during tissue dissection (Kasanga et al., 2024). The same assay procedure was done for the new results from the Pink1 cohort. For NfL, the R-PLEX Human NfL assay was used (cat# K1517XR-2; Meso Scale Discovery, Rockville, MD, United States) and for GFAP, the R-PLEX Human GFAP Assay (cat# K1511MR-2; Meso Scale Discovery, Rockville, MD, United States), respectively. Species and matrix compatibility were validated using linearity and spike recovery. UCHL-1 assays used an ELISA kit specific for rat, in a detection range of 62.5- 4000 pg/ml, sensitivity 37.5 pg/ml) (Biomatik, cat # EKE62409), and was validated for sample compatibility and accurate analyte measurement. Serum from all experimental groups were run in duplicate and incubated overnight. All other directions were followed according to the manufacturer’s protocol. Plates were read on MESO QuickPlex SQ 120 and analyzed using the DISCOVERY WORKBENCH analysis software (version 4.0).

### Statistical Analysis

Statistical analysis was performed using GraphPad Prism (Version 10). To ensure that the biomarker profiles reflected the degree of TH protein loss seen in human PD at motor symptom onset (Loss of TH at >80% in striatum and >50% in SN), two criterion were required for inclusion into statistical evaluation of the biomarkers in the 6-OHDA study; 1) the rats must have exhibited motor impairment by the 7^th^ day after lesion by evidence of ≥ 30% loss of forelimb use, and 2) TH protein loss must have exceeded 50% in the SN or 80% in the striatum. Of the 11 rats in the 7 days post-lesion group, 8 satisfied this criterion. Of the 13 rats in the 28 days post-lesion group, 9 rats satisfied this criterion. For the 6-OHDA cohort (which included the sham-operation control group), a repeated measures 3 way ANOVA was used to determine if the independent variables of sham vs lesion, side of treatment (sham-operation vs intact in the sham group and lesion vs contralateral to lesion in the 6-OHDA group), and days past treatment (7 or 28 days after sham operation or lesion induction) were significant or if there was interactions between two or all 3 independent variables. Significant results with independent variables alone or by interaction were then followed with 2-way repeated measures ANOVA to independently evaluate the impact of sham-operation vs 6-OHDA lesion on biomarker expression. In the Pink1 KO cohort, a 2-way ANOVA (not repeated measures) was used to determine genotype, aging, or interactions on biomarker results, followed by unpaired t-tests with Welch’s Correction if differences in variance between groups existed (as needed), if significance was detected for each variable or interaction between them.

To evaluate the relationship between biomarker expression levels and DA or TH protein expression levels, a Pearson or Spearman correlation was used depending on whether the results passed or did not pass, respectively, normality tests. The Grubb’s test was used to identify any outliers with alpha= 0.05, based on the *n* being evaluated.

## RESULTS

### Baseline expression of biomarkers in striatum and SN and tissue injury impact

The biomarker expression profile from the sham group represents a brain tissue injury response to differentiate against the biomarker profile associated with progressive nigrostriatal neuron loss after 6-OHDA. It also served as a relevant control to the side contralateral to lesion to ascertain whether changes in biomarker expression profile therein were influenced by the progressive loss of nigrostriatal neurons in the opposite hemisphere.

Sham-operation did not affect expression of UCHL-1 in the SN or striatum at either time point post-surgery, although there was notable variability in expression in both areas (Suppl Fig S1). GFAP expression was affected in the SN, but not striatum (Suppl Fig S2), with an unexpected decrease in expression in the intact side only at the latter time point after surgery. NfL expression was unaffected by sham-operation in the SN, but there was a transient increase in expression 7 days after the sham-operation in the striatum (Suppl Fig S3). Taken together, sham-operation transiently affected expression of GFAP in the SN and NfL in the striatum. These outcomes were accommodated for in the statistical evaluation of the impact of nigrostriatal neuron loss and pathology on biomarker expression in the 6-OHDA-lesioned rats.

### Loss of TH and DA neurons after 6-OHDA or aging in Pink1 KO

Our 6-OHDA lesion model produces differential TH protein loss between the striatum and SN, with maximal TH protein loss by 7 days only in the striatum (Kasanga et al., 2023). Striatal tissue collected from rats where cell loss was previously assessed (Kasanga et al., 2023) confirmed this severe TH loss by day 7 (lesion, F(1,9)=107, *p*<0.0001), with time after lesion having no significant effect on loss severity (days post lesion x lesion, F(1,9)=1.83, *p=*0.21) (Fig. 2A). However, despite the severity of TH protein loss in the striatum, in the SN, we previously reported that both TH protein loss (lesion, F(1,15)=62, *p*<0.0001; time after lesion F(1,6)=17.9, *p=*0.0006) (Fig. 2B) and TH+ cell loss (Fig. 2C) were progressive in the same time frame, with TH protein loss occurring at a faster rate than TH+ cell loss (Kasanga et al., 2023). Thus, 7 days after 6-OHDA lesion, striatal TH protein loss was 92%, nigral TH protein loss was 60%, and TH+ cell loss even less ∼23% (Fig. 2C). However, by day 28, both TH protein and cell loss in the SN approached similar severity of loss (∼85%) similar (Kasanga et al., 2023). In figure 2, we show two 6-OHDA studies (Kasanga et al., 2023; 2024) had a similar magnitude of TH+ cell loss, with new results showing comparable level of TH+ cell loss by day 28/29 after 6-OHDA lesion induction (Fig. 2D). As new (UCHL-1) and previously reported (GFAP and NfL) BB biomarker results were obtained from Kasanga et al., 2024, this new result demonstrates consistency in TH+ cell loss between two studies using 6-OHDA. Finally, whereas this model does not produce TH loss in striatum contralateral to lesion, there is TH loss in the contralateral SN between 20 and 40% by day 28 (Kasanga et al., 2023; Galfano et al., 2026). Thus, these disparities in TH protein expression and TH+ cell loss enabled determination of whether the severity of TH protein loss is related to biomarker expression levels in either region, and whether the BB biomarker profile correlated to biomarker expression profile in striatum or SN when TH+ cell loss was severe at 28 days post-6-OHDA.

In the Pink1 KO, we previously reported (Soto et al., 2024a) that aging accelerated TH protein loss and DA loss exclusively in the SN (Fig. 2F), without any loss of TH in the striatum (Fig. 2E). Notably, this decrease in the SN was exclusive to the Pink1 KO, as no loss was observed in the wild-type strain- and aged-matched control.

These similarities and differences between the two PD models enabled us to examine whether TH protein loss in the striatum or SN, per se, was associated with changes in biomarker expression.

### UCHL1 in SN and Striatum

#### Progressive 6-OHDA lesion

In the striatum, 3-way ANOVA revealed a significant interaction between time after treatment x side of treatment (F(1,26)=4.65, *p=*0.041). In the lesioned cohort alone, neither lesion (F(1,15)=0.19, *p*=0.66), nor day after lesion (F(1,16)=0.00, *p*=0.94) affected expression. There was a significant interaction between day after lesion and lesion (F(1,15)=6.12, *p=*0.026). However, differences in expression levels were not significant when comparing days post-lesion or side of lesion (Fig. 3A,C).

**Figure 3.**
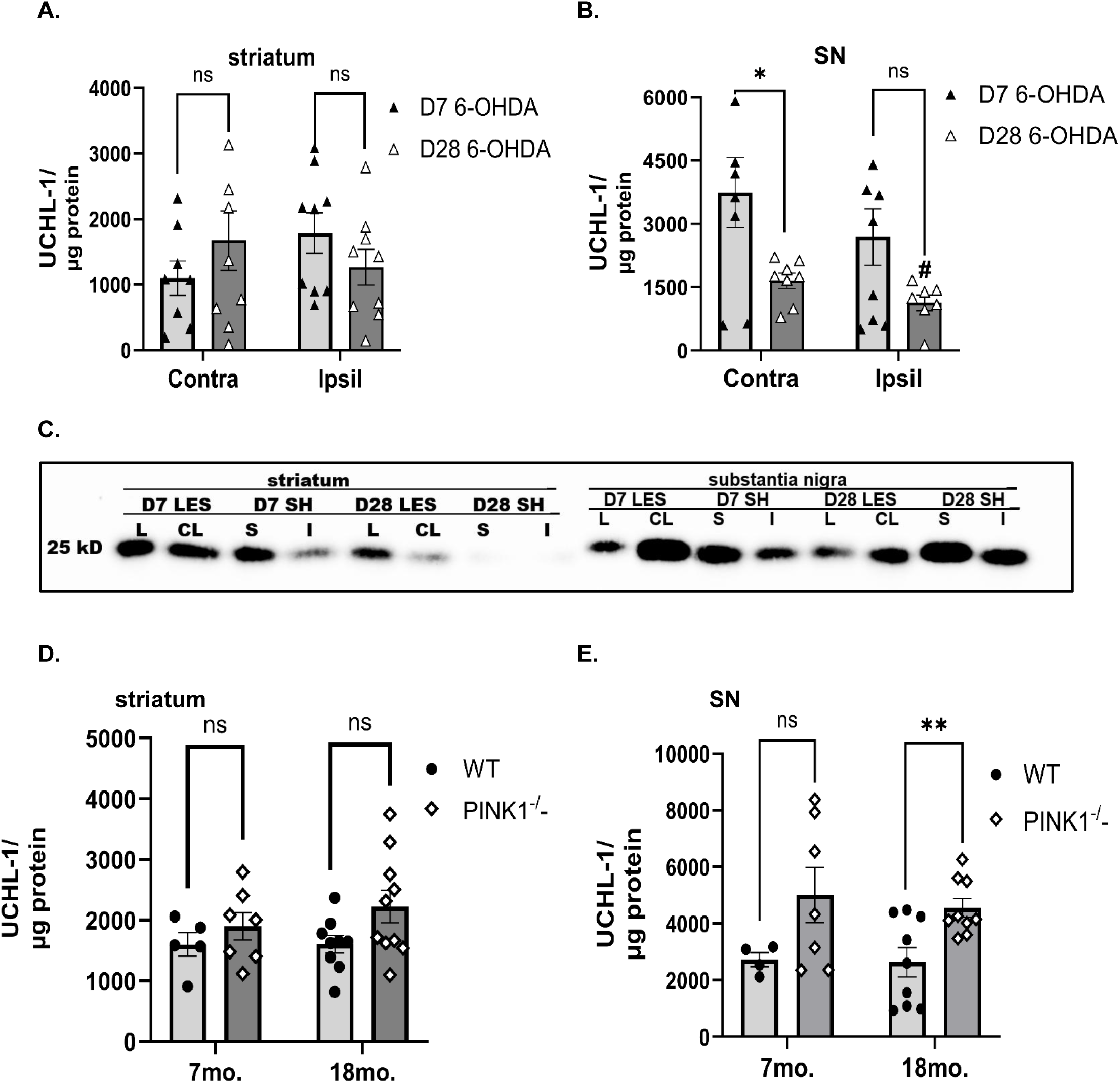
UCHL-1 expression during 6-OHDA-induced nigrostriatal neuron loss (A,B) at 7 and 28 days (D) post-lesion and in aging Pink1 KO (C,D). A. 6-OHDA, Striatum. Although there was a significant interaction between days post-lesion and side of lesion, UCHL-1 was not significantly different between the contralateral and ipsilateral sides or day after lesion. Ipsil v Contra. Day 7 (t=2.22, *p=*0.062), Day 28 (t=1.31, *p=*0.22). Day 7 v Day 28. Ipsil (t=1.27, *p=*0.22), Contra (t=1.05, *p=*0.31). **B. 6-OHDA, SN.** UCHL-1 expression decreased as nigrostriatal loss progressed after 6-OHDA-induced lesion. Ipsil v Contra. Day 7 (t=1.59, *p=*0.163), Day 28 (t=2.69, ^#^*p=*0.036). Day 7 v Day 28. Ipsil (t=2.00, *p=*0.065), Contra (t=2.48, \**p=*0.027). **C. Representative western blot images of UCHL-1 expression in striatum and SN across treatment groups in 6-OHDA and sham-operation cohorts.** Relative changes in UCHL-1 expression across treatment groups (6-OHDA lesioned (LES) vs. sham-operation (SH); Day past treatment (Day 7 (D7), Day 28 (D28), and side of treatment (ipsilateral to lesion (L), contralateral to lesion (CL), side of sham-operation (S), contralateral to sham or intact (I)) are depicted in this western blot image. **D. Pink 1 KO, striatum.** UCHL-1 expression was not affected in striatum either by genotype or age. **E. Pink 1 KO, SN.** UCHL-1 expression levels were higher in the KO vs WT at 7 months old, but this difference was not significant ((t=2.26, *p=*0.060) unlike that at 18 months old (t=3.31, \*\**p=*0.005).

In the SN, 3-way ANOVA revealed a significant interaction between time after treatment x lesion v sham (F(1,26)=4.51, *p=*0.044). In the lesioned cohort alone, UCHL-1 expression was decreased in the SN (side of lesion (F(1,13)=5.81, *p*=0.032); days post-lesion (F(1,15)=7.09, *p*=0.018), with lowest expression levels found both ipsilateral and contralateral to lesion by day 28 (Fig. 3B,C).

#### Pink 1 KO

In striatum (Fig. 3D), UCHL1 expression was unaffected in the Pink1 KO vs WT, with no significant genotype effect (F(1,27)= 3.77, *p=*0.063), aging effect (F(1,27)= 0.49, *p=*0.49), or genotype x aging interaction (F(1,27)= 0.45, *p=*0.51) found. Conversely in the SN, there was a highly significant effect of genotype on UCHL-1 expression in the SN (F(1,25)= 10.9, *p*= 0.003) (Fig. 3E), with significantly increased UCHL1 expression levels in 18 month-old Pink1 KO rats compared to age-matched WT controls. Age alone did not affect UCHL-1 expression (F(1,25)= 0.05, *p*=0.82), nor was there an age x genotype interaction (F(1,25)= 0.12, *p=*0.74).

#### UCHL-1 expression and relationship to DA tissue content

UCHL-1 expression was affected in opposite directions in the SN between the two PD models, with UCHL-1 decreasing in the 6-OHDA model and increasing in the aged Pink1 KO. As UCHL-1 is expressed in nigrostriatal neurons (Xilouri et al., 2012), this dichotomous result between PD models suggests that the frank 80% loss of nigrostriatal neurons in the 6-OHDA model contributed to UCHL-1 loss, masking an increase that may have occurred in remaining cells in response to cellular stressors, as occurs in the Pink1 KO. We found UCHL-1 had significant positive correlation to DA tissue content in both 6-OHDA (Fig. 4A) (contralateral and ipsilateral to lesion combined, r=0.587, *p=*0.0006, *n=*30) and in the Pink 1 KO cohort (Fig. 4B) (Pink 1 and WT combined, r=0.622, *p*=0.0002, *n=*30). The similar findings between the two models suggests UCHL-1 levels influence DA tissue levels.

**Figure 4.**
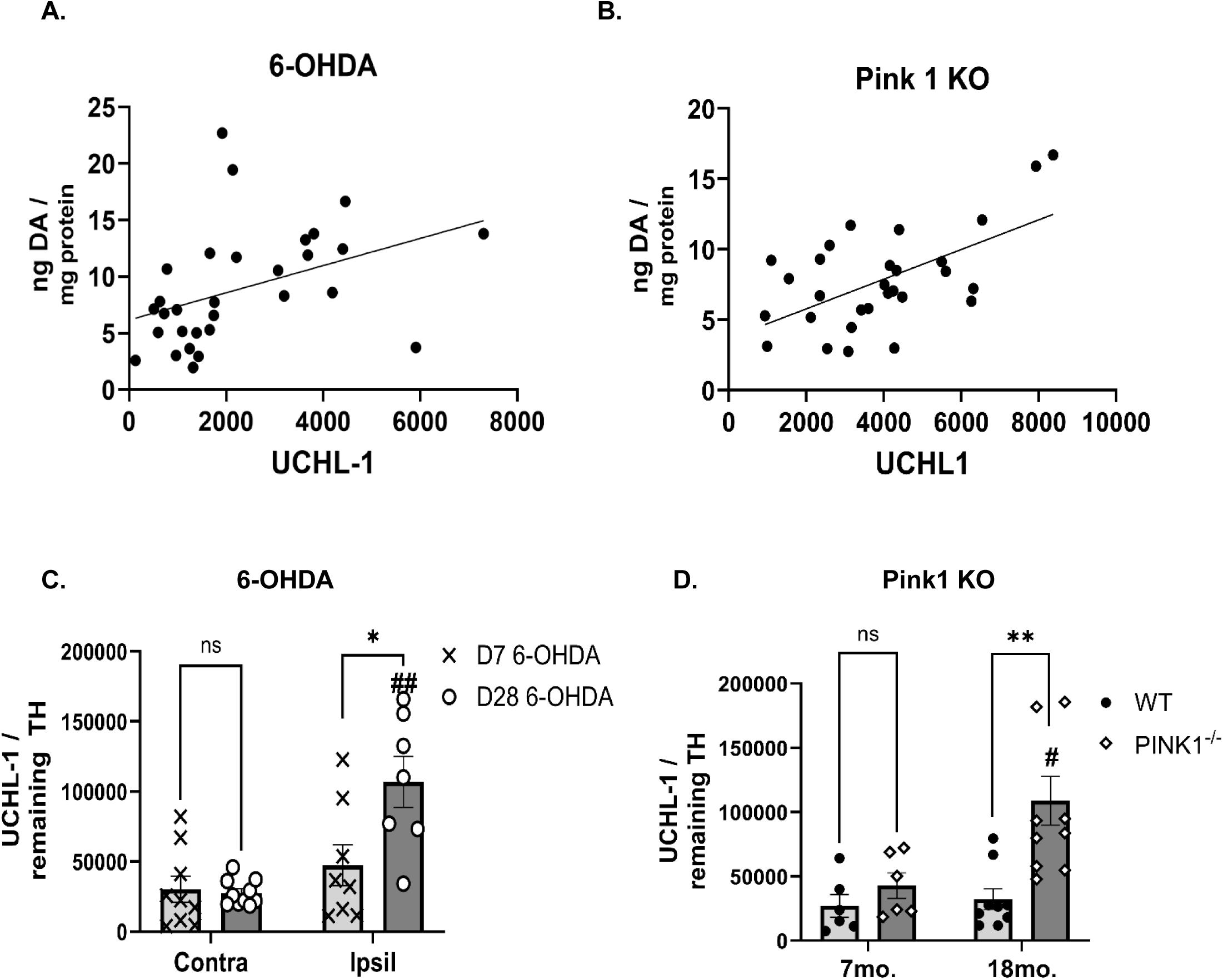
Relationship of UCHL-1 expression as a function of TH loss severity after 6-OHDA or aging in the Pink 1 KO. A. Nigral UCHL-1 expression vs. DA tissue content, 6-OHDA cohort. UCHL-1 protein expression had significant positive correlation with DA tissue levels in the lesioned and contralateral to lesioned SN (r=0.587, *p=*0.0006, XY pairs=30, Spearman correlation. **B. Nigral UCHL-1 expression vs. DA tissue content, Pink1 KO cohort.** UCHL-1 protein expression had significant positive correlation with DA tissue levels in the WT and KO genotypes combined (r=0.622 *p=*0.0002, XY pairs=30, Pearson correlation). **C. Nigral UCHL-1 expression against remaining TH protein, 6-OHDA cohort.** Time past lesion induction played a significant role in UCHL-1 expression against TH protein loss (F(1,30)=6.19, *p=* 0.019). Lesion alone had a significant effect on UCHL-1 expression (F(1,30)=17.8, *p*=0.0002). Despite 58% TH loss by day 7, levels of UCHL-1 were not significantly different in lesioned vs contralateral to lesion SN (t=1.35, *p*=0.22). Greater TH protein loss by day 28 after 6-OHDA was associated with increased UCHL-1 expression in SN ipsilateral to lesion vs respective contralateral SN (t=4.10, ^##^*p*=0.009) and greater levels on the lesioned SN at day 28 vs day 7 (t=2.67, \**p*=0.020). There was no difference in expression between day 7 and 28 in SN contralateral to lesion (t=0.24, *p*=0.81). **D. Nigral UCHL-1 expression against remaining TH protein, Pink1 KO cohort.** UCHL-1 expression increased in the SN of the aged Pink1 KO against the young Pink 1 KO (t=2.55, ^#^*p*=0.023), and vs the aged WT (t*=*3.58, \*\**p*=0.002).

Impairment of mitochondrial function and protein degradation occurs in both of these models (Elkon et al., 2001; Villeneuve et al., 2016; Gonçalves et al., 2019; Iravanour et al., 2021; Bouron et al., 2023). Thus, UCHL-1 levels were then normalized against remaining TH protein levels in both models to address whether changes in UCHL-1 expression were consistent as a function of mitochondrial impairment presumed to worsen as TH loss increased. In the 6-OHDA model, TH protein and cell loss is progressive in the SN between day 7 (58%) and day 28 (84%) (Kasanga et al., 2023), suggesting a greater level of mitochondrial impairment by day 28, as previously shown in this model (Kupsch et al., 2014). UCHL-1 expression increased as TH expression decreased to severe loss between day 7 and day 28 (F(1,30)=7.38, *p*=0.011) (Fig. 4C). Moreover, in the Pink1 KO, the levels of UCHL-1 per TH protein also increased in the Pink1 KO (F(1,27)=10.0, *p*=0.004), but only in the aged KO (F(1,27)=6.00, *p=*0.021); lesion X genotype (F(1,27)=4.33, *p=*0.047) (Fig. 4D). Together, these data indicate UCHL-1 expression increases in the SN, but not striatum, when cellular stress or mitochondrial dysfunction reaches a critical threshold in the nigrostriatal pathway.

### GFAP in SN and Striatum

#### Progressive 6-OHDA lesion

In the striatum, 3-way ANOVA revealed a sham vs lesion effect on GFAP expression (F(1,48)=5.39, *p*=0.025), with significant interaction between time after treatment x side of treatment (F(1,48)=7.42, *p=*0.009). In the lesioned cohort alone, there was a significant interaction between day after lesion and lesion (F(1,28)=6.77, *p=*0.015), with increased GFAP expression on day 7 ipsilateral vs contralateral to lesion, but not day 28 after 6-OHDA (Fig. 5A,C). Neither lesion (F(1,28)=0.01, *p*=0.94), nor day after lesion (F(1,28)=0.54, *p*=0.47) alone affected expression.

**Figure 5.**
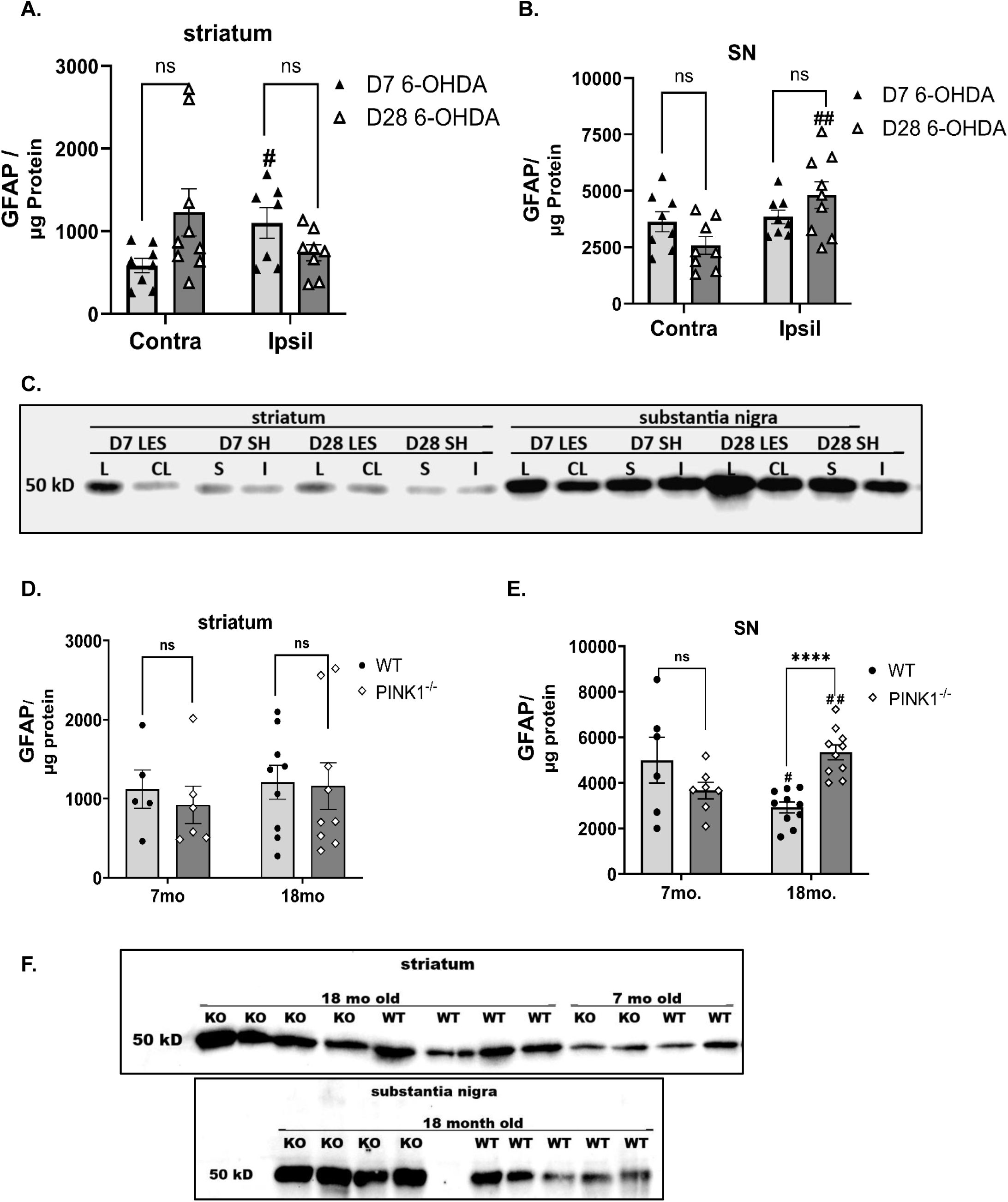
GFAP expression during 6-OHDA-induced nigrostriatal neuron loss (A,B) at 7 and 28 days (D) post-lesion and in aging Pink1 KO (C,D). **A. 6-OHDA, Striatum.** There was a significant interaction between days post-lesion and lesion, with increased GFAP expression (lesion v contra lesion) on day 7 (t=2.49, ^#^*p=*0.047), but not day 28 (t=1.01, *p=*0.35). Day 7 v Day 28. Ipsil (t=1.80, *p=*0.10), Contra (t=2.04, *p=*0.06). **B. 6-OHDA, SN.** GFAP expression increased as nigrostriatal loss progressed after 6-OHDA-induced lesion. Ipsil v Contra. Day 7 (t=0.76, *p=*0.46), Day 28 (t=3.07, ^##^*p=*0.008). Day 7 v Day 28. Ipsil (t=1.300, *p=*0.19), Contra (t=1.78, *p=*0.097). **C. Representative western blot images of GFAP expression in striatum and SN across treatment groups in 6-OHDA and sham-operation cohorts.** Relative changes in GFAP expression across treatment groups (6-OHDA lesioned (LES) vs. sham-operation (SH); Day past treatment (Day 7 (D7), Day 28 (D28), and side of treatment (ipsilateral to lesion (L), contralateral to lesion (CL), side of sham-operation (S), contralateral to sham or intact (I)) are depicted in this western blot image. **D. Pink 1 KO, striatum.** GFAP expression was not affected in striatum either by genotype or age. **E. Pink 1 KO, SN.** There was no significant difference in GFAP expression between WT and KO at 7 months old (t=1.33, *p=*0.21). At 18 months old, GFAP expression increased >50% in the KO vs WT (t=5.94 \*\*\*\**p<*0.0001) and was also greater vs the 7 month old KO (t=3.36, ^##^*p=*0.0043). The opposite effect on GFAP expression was observed in the WT (t=2.53, ^#^*p=*0.024). **F. Representative western blot images of GFAP expression in striatum and SN in Pink1 KO vs. age-matched wild-type (WT).**

In the SN, 3-way ANOVA revealed a significant effect of treatment side (F(1,22)=22.1, *p=*0.0001) and interaction between treatment side (F(1,22)=7.62, *p=*0.011). In the lesioned cohort alone, GFAP expression increased in the SN (lesion (F(1,14)=12.1, *p*=0.004) and there was an interaction between lesion day after lesion and lesion (F(1,14)=8.33, *p*=0.012), with increased GFAP expression ipsilateral vs contralateral to lesion on day 28 (Fig. 5B,C).

#### Pink 1 KO

In the striatum (Fig. 5D,F), no significant genotype effect (F(1,25)= 0.02, ns), aging effect (F(1,25)= 0.34, ns), or genotype x aging interaction (F(1,25)= 0.37, ns) was observed, on GFAP expression, with higher noted variability. In the SN, there was a significant genotype effect (F(1,25)= 11.3, *p*= 0.003), but no effect of aging (F(1,285= 0.12, p= 0.73) or genotype x aging interaction (F(1,25)= 0.04, p= 0.83) on GFAP expression (Fig. 5E,F). This was attributed to increased GFAP levels in Pink1 KO rats from 7 to 18 months, and higher expression in 18-month-old Pink1 KO rats compared to age-matched WT rats.

#### GFAP expression and relationship to DA tissue content

As GFAP expression was increased in the SN, but not striatum, in the latter stages of TH protein loss in the SN in both PD models, we evaluated whether GFAP expression would co-vary with DA tissue levels or TH protein. We found different outcomes between the two models, with no correlation of GFAP with nigral DA tissue content in the 6-OHDA model (r= -0.105, *p=*0.56, *n=*33) (Fig. 6A), but a negative correlation between GFAP and DA tissue content in the Pink 1 KO r= - 0.406, *p=*0.019, *n=*33) (Fig. 6B), indicating that GFAP expression increased as DA tissue content decreased in the SN of the aged Pink1 KO.

**Figure 6.**
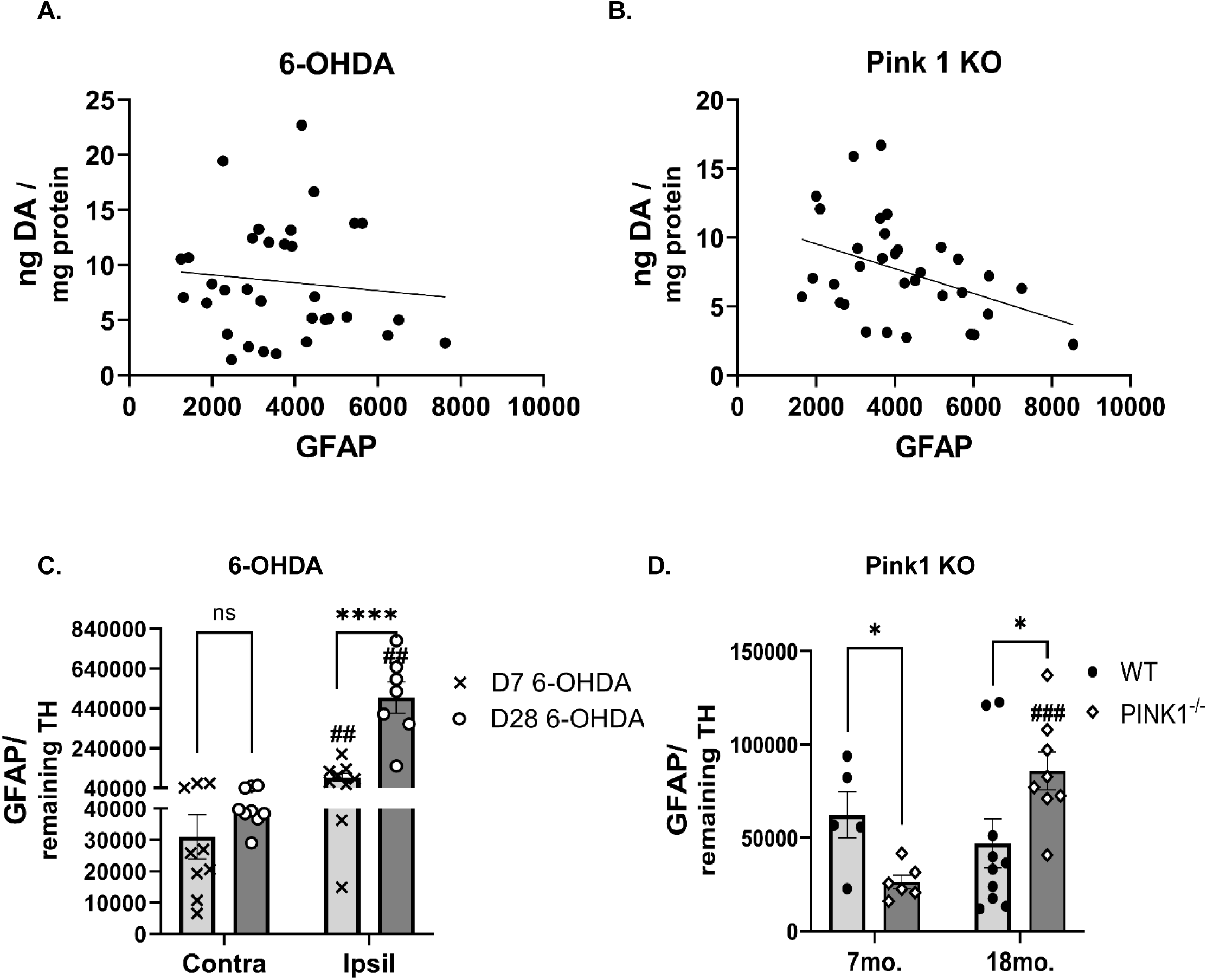
Relationship of GFAP expression as a function of TH loss severity after 6-OHDA or aging in the Pink 1 KO. A. Nigral GFAP expression vs DA tissue content, 6-OHDA cohort. GFAP protein expression had no correlation with DA tissue levels in the lesioned and contralateral to lesioned SN. **B. Nigral GFAP expression vs. DA tissue content, Pink1 KO cohort.** GFAP protein expression had significant negative correlation with DA tissue levels in the WT and KO genotypes combined (r= -0.622 *p=*0.0002, XY pairs=30, Pearson correlation. **C. Nigral GFAP expression against remaining TH protein, 6-OHDA cohort.** Time past lesion induction played a significant role in GFAP expression against TH protein loss (F(1,16)=36.2, *p<* 0.0001). Lesion alone had a significant effect on GFAP expression (F(1,14)=59, *p*=0.0002). By 7 days after 6-OHDA lesion, GFAP per remaining TH protein increased (t=4.25, ^##^*p*=0.003) and greater levels on the lesioned SN at day 28 vs day 7 (t=5.68,^##^*p*=0.001). There was no difference in expression between day 7 and 28 in SN contralateral to lesion (t=1.28, *p*=0.21), but GFAP expression against remaining TH increased substantially between day 7 and day 28 (t=5.58, \*\*\*\**p*<0.0001). **D. Nigral GFAP expression against remaining TH protein, Pink1 KO cohort.** GFAP expression against TH protein increased in the SN of the aged Pink1 KO against the young Pink 1 KO (t=4.85, ^###^*p*=0.0004), and vs the aged WT (t*=*2.25, \**p*=0.039). The young cohort showed the less GFAP/TH in the KO (t=3.04, \**p=*0.014).

Astrocytes may increase GFAP expression in response to nigrostriatal neuron loss or loss of function (Kasanga et al, 2023b). With the progressive loss of TH in the 6-OHDA model (day 7 (58%) vs day 28 (84%)), GFAP expression against remaining TH protein was increased both early and late after 6-OHDA lesion (lesion, F(1,14)=59.0, *p*<0.0001); days past lesion F(1,16)=36.2, *p*<0.0001); lesion X days past lesion F(1,14)=34.0, *p*<0.0001), with GFAP expression progressively increasing as TH loss increased in the SN between day 7 and day 28 (Fig. 6C). In the Pink1 KO, the levels of GFAP per TH protein increased in the Pink1 KO as a function of age (age X genotype) (F(1,25)=9.59, *p*=0.005), age (F(1,25)=3.37, *p=*0.078) (Fig. 6D). Together, these data suggest GFAP expression increases as nigrostriatal neuron loss progresses after 6-OHDA lesion or during aging as mitochondrial impairment and TH loss occurs in the Pink1 KO.

### NfL in SN and Striatum

#### Progressive 6-OHDA lesion

In the striatum, 3-way ANOVA did not reveal any significant effect of lesion vs sham (F(1,27)=0.15, *p*=0.70), or any significant interaction with time after treatment (F(1,27)=1.38, *p*=0.25) or side of treatment (F(1,20)=1.07, *p=*0.31). In the lesioned cohort alone, no significant interactions or main effects were noted; lesion (F(1,11)=0.29, *p=*0.60), lesion x days past lesion (F(1,11)=0.08, *p=*0.78) (Fig. 7A,C).

**Figure 7.**
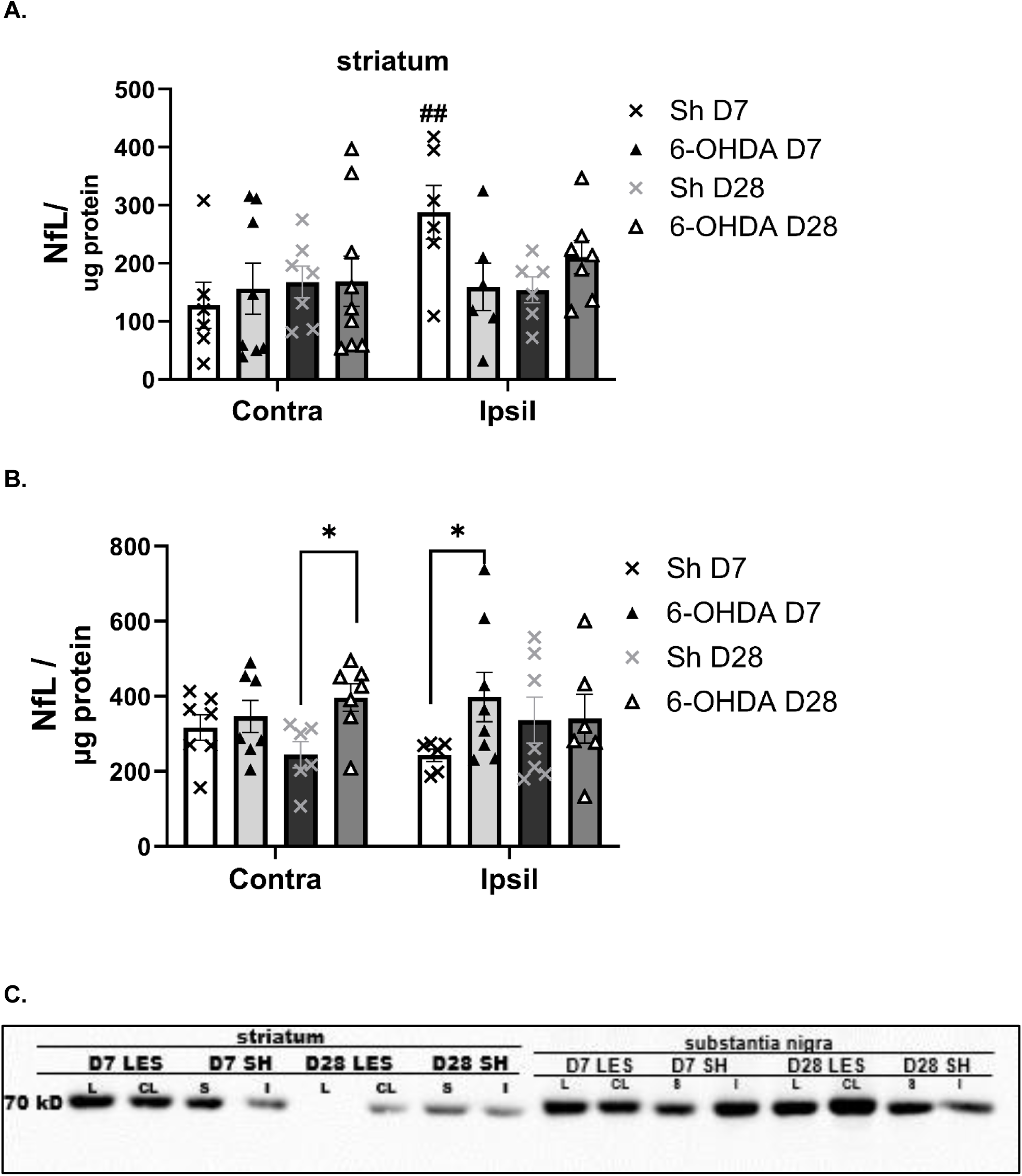
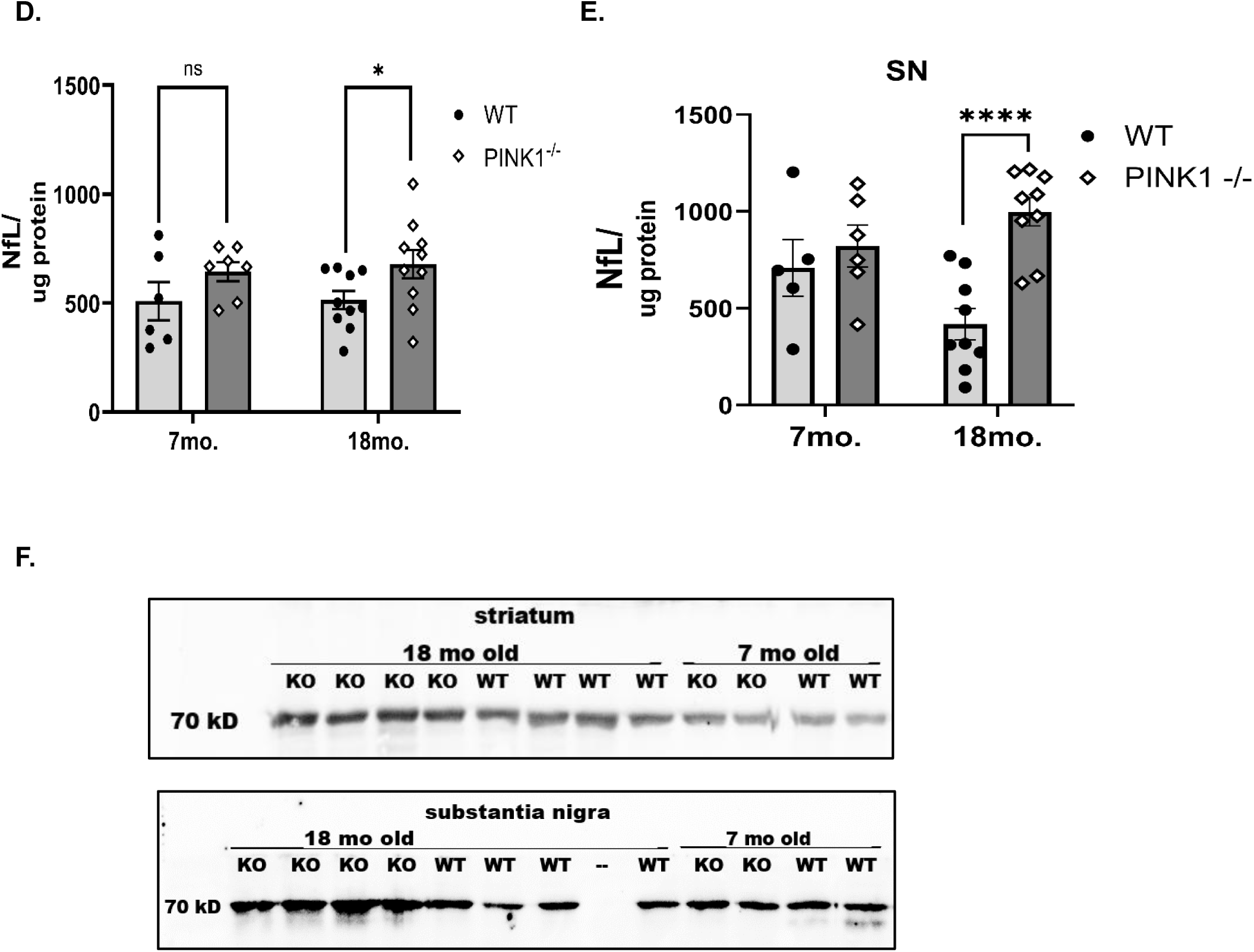
NfL expression during 6-OHDA-induced nigrostriatal neuron loss (A,B) at 7 and 28 days (D) post-lesion and in aging Pink1 KO (C,D). A. 6-OHDA, Striatum. Although the sham-operation transiently increased NfL expression, lesion had no effect on either side (Contra or Ipsil) of 6-OHDA lesion. **B. 6-OHDA, SN.** NfL expression increased bilaterally early and later after 6-OHDA lesion induction increased as nigrostriatal loss progressed after 6-OHDA-induced lesion. Sham v lesion; Ipsil, Day 7 (t=2.30, \**p=*0.051), Day 28 (t=0.04, *p=*0.96); Contra, Day 7 (t=0.540, *p=*0.60), Day 28 (t=3.00, \**p=*0.012). **C. Representative western blot images of NfL expression in striatum and SN across treatment groups in 6-OHDA and sham-operation cohorts.** Relative changes in NfL expression across treatment groups (6-OHDA lesioned (LES) vs. sham-operation (SH); Day past treatment (Day 7 (D7), Day 28 (D28), and side of treatment (ipsilateral to lesion (L), contralateral to lesion (CL), side of sham-operation (S), contralateral to sham or intact (I)) are depicted in this western blot image. **D. Pink 1 KO, striatum.** NfL expression was not affected in striatum either by genotype or age. **E. Pink 1 KO, SN.** There was no significant difference in NfL expression between WT and KO at 7 months old (t=1.33, *p=*0.21). At 18 months old, NfL expression increased >50% in the KO vs WT (t=5.94 \*\*\*\**p<*0.0001) and was also greater vs the 7 month old KO (t=3.36, ^##^*p=*0.0043). The opposite effect on GFAP expression was observed in the WT (t=2.53, ^#^*p=*0.024). **F. Representative western blot images of NfL expression in striatum and SN in Pink1 KO vs. age-matched wild-type (WT).**

In the SN, 3-way ANOVA revealed a significant effect of lesion vs sham (F(1,27)=5.36 *p=*0.028), with no other significant interactions with time after treatment (F(1,27)=0.025, *p=*0.87) or side of treatment (F(1,19)=0.02, *p=*0.88). However, in the lesioned cohort alone, NfL expression was not significantly different as a function of side of lesion, nor was there an interaction with days post lesion (F(1,9)=0.98, *p*=0.35). Notably, this lack of difference within the lesion group alone was due to a bilateral increase in NfL that occurred in both the ipsilateral and contralateral side to lesion, as indicated by the 3 way ANOVA result (Fig. 7B,C). At day 7, NfL levels increased in the SN ipsilateral to lesion vs. sham-op levels and at day 28, NfL levels increased in the SN contralateral to lesion.

#### Pink 1 KO

In striatum, NfL expression was moderately greater in the Pink1 KO (Fig. 7D,F) (genotype (F(1,29)= 6.11, *p*= 0.020)), with this difference being significant in the aged cohort, although there was not a significant genotype x aging interaction (F(1,29)= 0.06, *p*= 0.81). In the SN, NfL expression was significantly influenced by genotype (F(1,25) 12.5, *p=*0.002), and this difference was further increased by aging (genotype x aging interaction (F(1,25)= 5.67,*p* =0.025) (Fig. 7E,F) NfL levels were substantially greater (more than 2-fold) in the aged Pink1 KO compared to the aged-matched WT.

#### NfL expression and relationship to DA tissue content

NfL expression was also increased in the SN, but not striatum, in both PD models. We evaluated whether this nigra-specific increase had a potential influence on DA tissue levels and found that in both models, NfL expression had positive correlation with nigral DA tissue content in the 6-OHDA model (r= 0.410, *p=*0.034) (Fig. 8A), and in the Pink 1 KO r= - 0.368, *p=*0.038) (Fig. 8B), indicating that the loss of DA seen in these models portends NfL protein loss in the CNS. This makes it possible that if NfL is no longer associated with DA neuropil due to its loss, the NfL is solubilized in extracellular fluid eventually making its way into the systemic circulation. With the progressive loss of TH in the 6-OHDA model, NfL expression against remaining TH protein progressively increased both early and late after 6-OHDA lesion (lesion, F(1,10)=22.1, *p*=0.0008); days past lesion F(1,15)=13.7, *p*=0.002); lesion X days past lesion F(1,10)=8.2, *p*=0.017) (Fig. 8C). Notably, NfL also increased in the side contralateral to lesion by Day 28. In the Pink1 KO, the levels of NfL per TH protein increased as a function of age (F(1,25)=12.9, *p=*0.001), genotype (F(1,25)=10.1, *p*=0.004), and interaction between genotype and age (age X genotype) (F(1,25)=13.1, *p*=0.001) (Fig. 8C). NfL expression was substantially greater in the aged KO cohort compared to both the young KO and the aged WT KO (Fig 8D). Together, these data suggest that NfL expression increases in the 6-OHDA model (after lesion) with progressive loss of nigrostriatal neurons, and in the Pink1 KO model with age, mitochondrial impairment, and TH loss.

**Figure 8.**
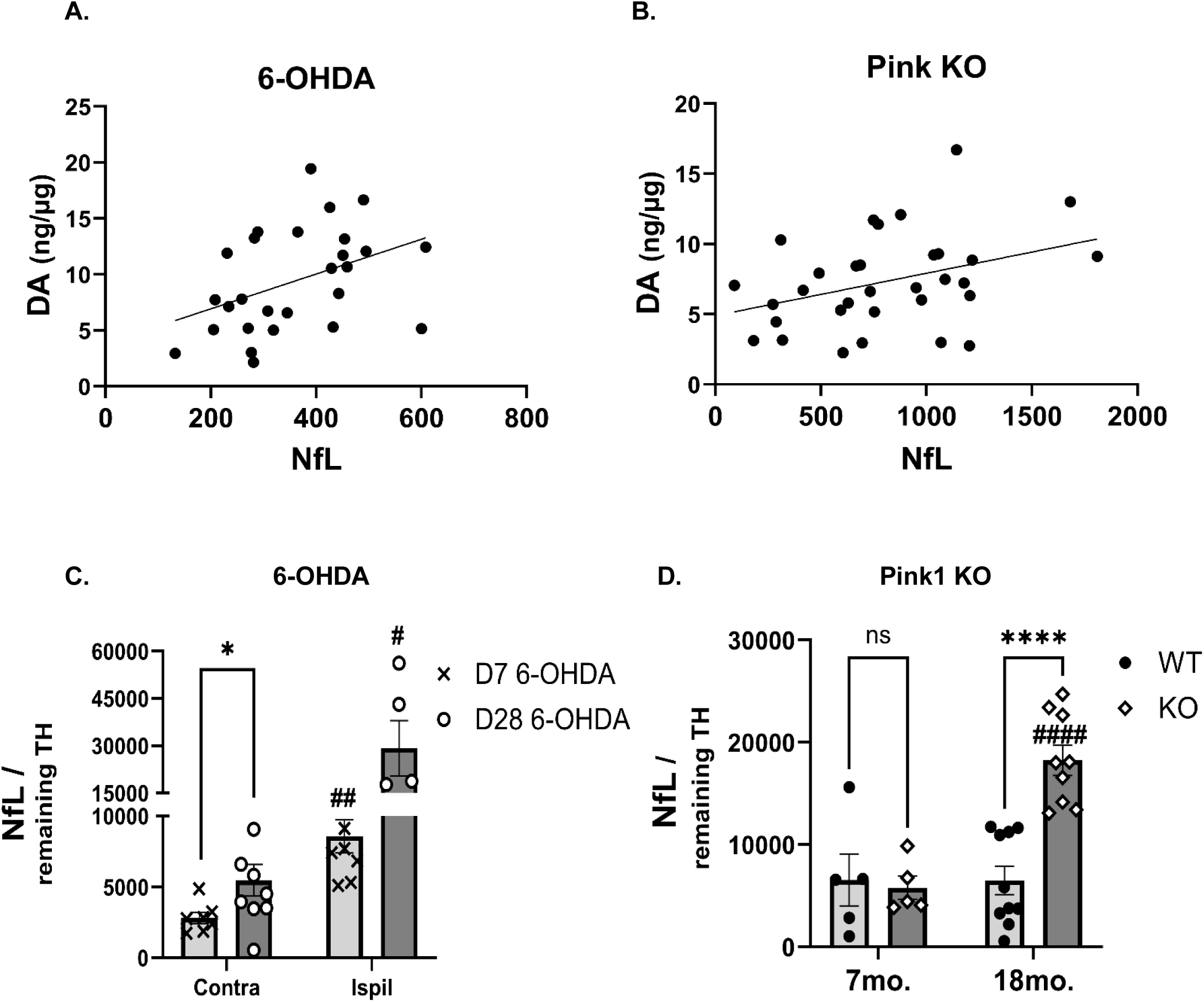
Relationship of NfL expression as a function of TH loss severity after 6-OHDA or aging in the Pink 1 KO. A. Nigral NfL expression vs. DA tissue content, 6-OHDA cohort. NfL protein expression had significant positive correlation with DA tissue levels in the lesioned and contralateral to lesioned SN (r=0.410, *p=*0.034, XY pairs=27), Pearson correlation. **B. Nigral NfL expression vs. DA tissue content, Pink1 KO cohort.** NfL protein expression had significant positive correlation with DA tissue levels in the WT and KO genotypes combined (r=0.368 *p=*0.038, XY pairs=32, Pearson correlation. **C. Nigral NfL expression against remaining TH protein, 6-OHDA cohort.** NfL expression increased on side of lesion at both time points after lesion induction, with increases seen by day 7 (Contra v Ipsil) (t=4.71, ^##^*p*=0.005), and further increasing at day 28 (t=3.50, ^#^*p* =0.025). The increase was greater by day 28 vs day 7, but not significant (t=2.39, *p*=0.077), likely due to the increase seen on the side contralateral to lesion by day 28 (Contra; day 7 v day 28 (t=2.63, \**p*=0.028). **D. Nigral NfL expression against remaining TH protein, Pink1 KO cohort.** NfL expression increased in the SN of the aged Pink1 KO against the young Pink 1 KO (t=5.70, ^####^*p*<0.0001), and vs the aged WT (t*=*5.77, \*\*\*\**p*<0.0001).

#### Relative abundance of UCHL-1, GFAP, and NfL in serum

UCHL-1 levels in the serum increased ∼24% in the 6-OHDA-lesioned rats by 28 days after lesion induction versus sham-op group (Fig. 9A). In the Pink1 KO, there was a modest, but significant increase (∼22%), in UCHL-1 in the aged vs young KO rat (age X genotype) (F(1,25)=4.9 *p*=0.037), without any overall effect of age (F(1,25)=0.06, *p=*0.81), or genotype (F(1,25)=0.00, *p*=0.98) (Fig. 8B). Thus, genotype alone was not a significant influence on UCHL-1 expression (Fig. 9C).

**Figure 9.**
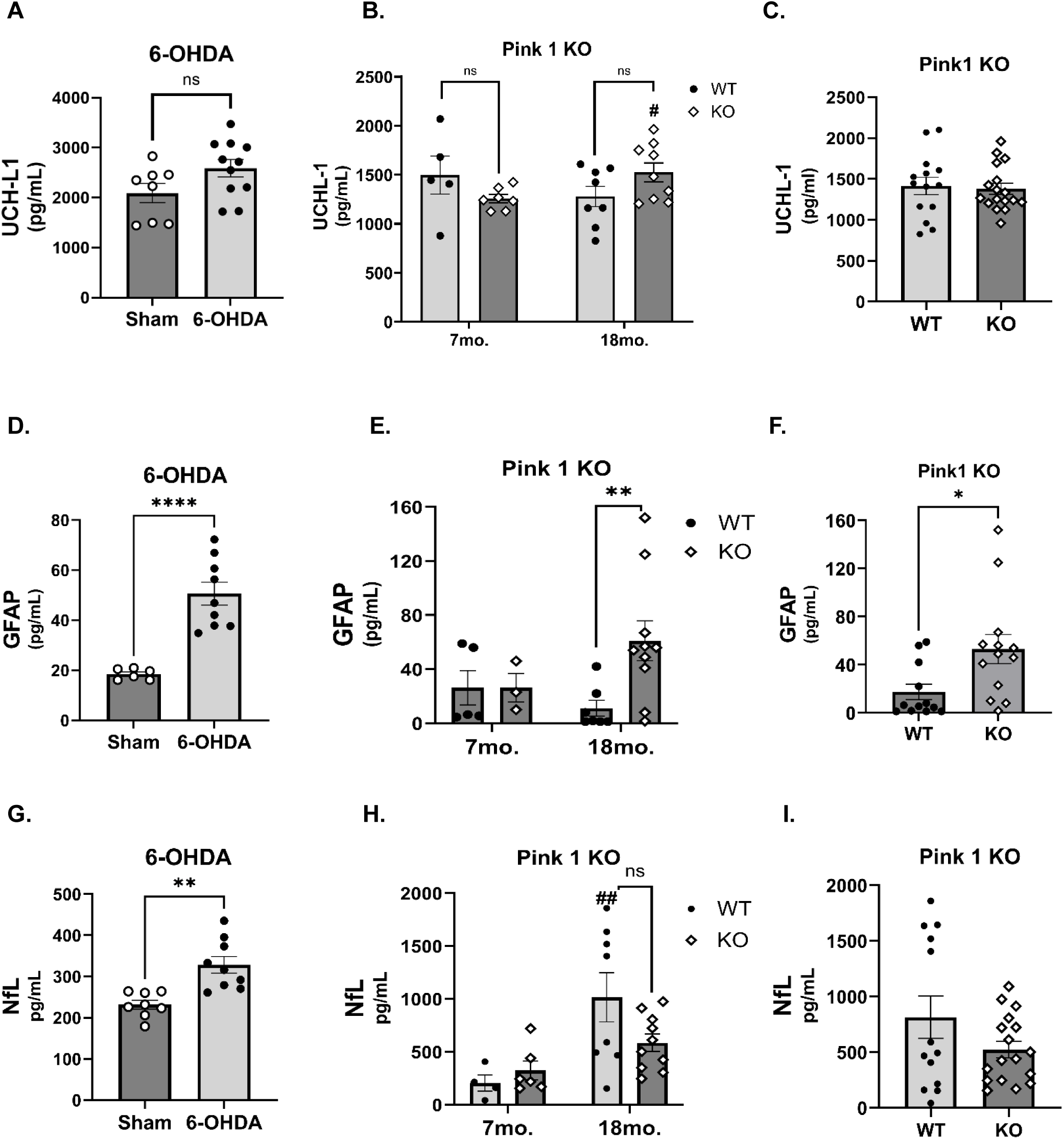
BB biomarker expression profile in serum 4 weeks after 6-OHDA lesion and genotype or age differences in Pink1 KO. Blood collected from trunk during tissue dissection was processed into serum and analyzed for differences in biomarker levels against rats receiving a sham-operation. **A. UCHL-1, 6-OHDA.** UCHL-1 levels were 1.25-fold greater in the 6-OHDA lesion group (t=1.92, *p*=0.071). **B. UCHL-1, Pink1 age x genotype.** There was a significant interaction between age and genotype, with an increase in UCHL-1 serum levels in the aged vs young Pink1 KO (t=2.56, *p*=0.027). **C. UCHL-1, genotype only.** There was no genotype difference when combining both age groups (t=0.29, *p*=0.78). **D. GFAP, 6-OHDA.** GFAP levels were more than 2-fold greater in the 6-OHDA lesioned (50.6 pg/ml) vs sham-operated rats (18.4 pg/ml) (t=6.83, \*\*\*\**p*<0.0001) (adapted from Kasanga et al., 2024). **E. GFAP, Pink1 age x genotype.** GFAP serum levels in the aged cohort were greater in the Pink1 KO (t=3.2, \*\**p*=0.008). **F. GFAP, genotype only.** The Pink1 KO had overall greater serum GFAP levels vs WT (t=2.6, *\*p*=0.018). **G. NfL, 6-OHDA.** NfL levels were more than 1.4-fold greater in the 6-OHDA lesioned (328 pg/ml) vs sham-operated rats (232 pg/ml) (t=4.05, \*\**p=*0.001) (adapted from Kasanga et al., 2024). **H. NfL, Pink1 age x genotype.** NfL serum levels in the aged cohort vs young cohort were greater in the WT (t=3.3, \*\**p*=0.010) and in the KO (t=2.1, *p*=0.059). **C. NfL, genotype only.** Genotype was not a factor affecting serum NfL (t=1.6, *p*=0.126).

With severe TH loss in the SN, 28 days after 6-OHDA lesion induction, serum levels of GFAP increased (Fig. 9D) (Kasanga et al., 2024), again notably greater than that associated with sham-operation, pointing to strong alignment with nigrostriatal neuron loss seen in this model (Kasanga et al., 2023a). In the Pink1 KO, we did not observe a difference in age (F(1,21)=0.42, *p=*0.53), but trends toward significance in genotype (F(1,21)=2.7, *p*=0.11), and interaction between age X genotype (F(1,21)=2.7, *p*=0.11). We noted wide variability in GFAP levels in the WT and KO groups, accounting for 3 and 2 outliers, respectively. There was a significantly greater level of GFAP in the aged Pink1 KO vs age-matched WT (Fig. 9E). Due to wide variability and relatively low n in the young groups, we combined the age groups to compare genotype, and found a significantly greater level of GFAP in serum of the Pink1 KO (Fig. 9F).

Serum levels of NfL increased with severe TH loss in the SN, 28 days after 6-OHDA lesion induction (Fig. 9G) (Kasanga et al., 2024). In the Pink1 KO, there was a strong influence of age on NfL levels (F(1,24)=10.9, *p=*0.003), but no effect of genotype (F(1,24)=0.92, *p*=0.35), but trend toward significance in interaction between age X genotype (F(1,24)=2.87, *p*=0.10) (Fig. 9H). There was also variability in NfL levels in the WT and KO groups, accounting for 3 and 1 outliers, respectively. NfL levels were significantly greater in the aged WT and KO, compared to their genotyped-match of younger age (Fig. 9H). Genotype was not a factor in the differences in serum NfL levels (Fig. 9I).

In summary, the moderate and variable increase in UCHL-1, and robust increase in GFAP and NfL (first reported in Kasanga et al., 2024), may be indicative of severe TH loss in the SN. In the case of the Pink1 KO wherein TH loss in SN occurred with aging, but not between genotypes, it appears that TH loss is associated with changes in UCHL-1 expression seen in aging, whereas for GFAP, genotype was associated with increased serum levels and not necessarily driven by aging. On the other hand, differences in NfL levels were driven only by aging in the Pink1 cohort.

#### Relative abundance of biomarkers between striatum and SN

To further ascertain how serum biomarker profile in the nigrostriatal pathway contributes to the expression profile in blood, we compared the relative expression levels of UCHL-1, GFAP, and NfL between the striatum and SN, using the sham-op cohort. Levels were greater in the SN than in striatum for all 3 biomarkers (Fig. 10 A-C), which would suggest that contribution of these biomarkers from these two regions would be greater from the SN than the striatum, particularly GFAP, which has ∼10-fold greater expression in the SN.

**Figure 10.**
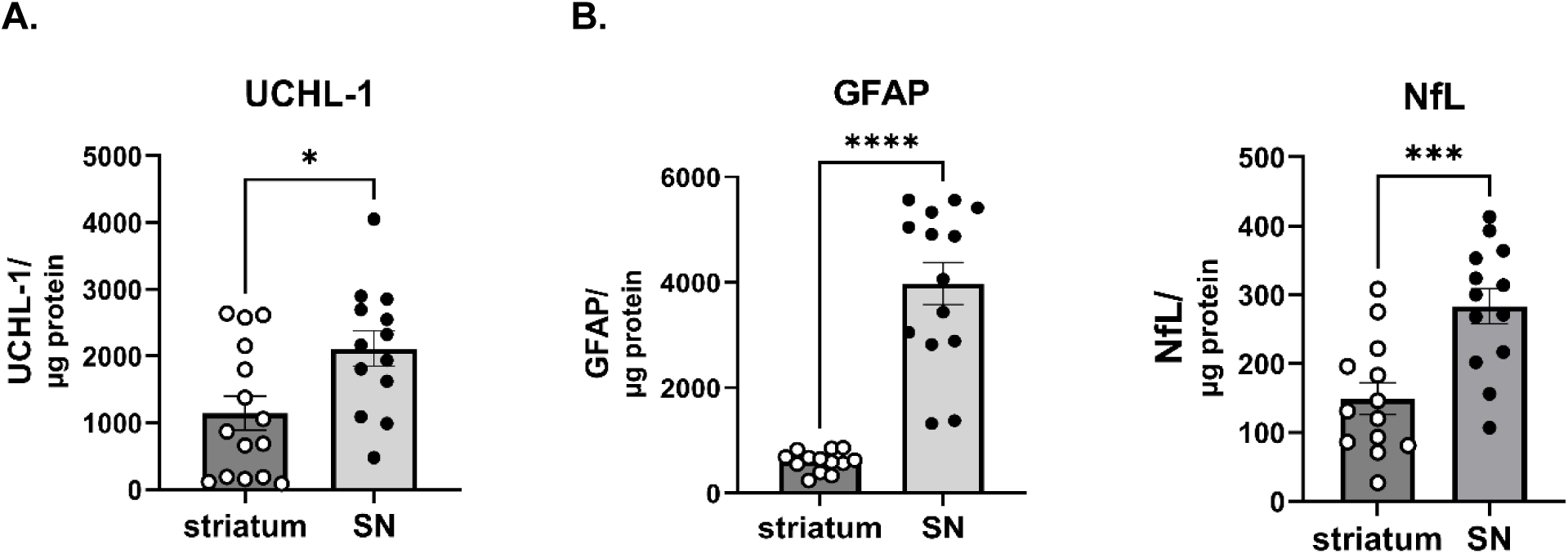
Regional differences in biomarker expression in nigrostriatal pathway; relative abundance of UCHL-1, GFAP, and NfL in striatum vs. SN. Striatum and SN tissue from intact (no-operation) hemisphere of the sham-op group from both the 7 and 28 day cohorts was used to compare relative abundance of biomarkers in the nigrostriatal pathway or allied tissue**. A. UCHL-1.** UCHL-1 expression was significantly greater in the SN vs striatum (t=2.64, \**p*=0.014). **B. GFAP.** GFAP expression was substantially greater in the the SN vs striatum (t=8.33, \*\*\*\**p*<0.0001). **C. NfL.** NfL expression was substantially greater in the SN vs striatum (t=3.89, \*\*\**p*=0.0007).

## DISCUSSION

Although BB biomarkers have become increasingly incorporated into characterizing PD severity, there is a significant knowledge gap in establishing their biological relevance to nigrostriatal pathology. Our preclinical study provides critical data to help close this gap by demonstrating changes in BB biomarker expression are in accord with changes in expression in the SN, and critically, correspond to decreased DA signaling therein in two rat PD models; the 6-OHDA unilateral hemi-lesion model and the Pink1 KO. We point out that these models represent different disease-relevant neurobiological backgrounds on different time scales, with the apparent key factor being the loss of TH protein and DA in the SN (Kasanga et al., 2023a; Soto et al., 2024a). The common neurobiological stressor between models is mitochondrial impairment (Kupsch et al., 2014; Villeneuve et al., 2016), which enabled us to determine if three BB biomarkers, previously shown to change in serum of PD patients, would change in expression in disease-relevant tissue. Specifically, the similarity in DA and TH protein loss in the SN and difference in severity of DA and TH protein loss in striatum between these models afforded insight into whether the increases in UCHL-1, GFAP, and NfL, shown to occur in PD patients, may be traced back to differences in their expression in these nigrostriatal neuron compartments (Bäckström et al., 2020; Buhmann eta al., 2023; Dong et al., 2023; Huang et al., 2022; Lin et al., 2023; Liu et al., 2022; Mollenhauer et al., 2020; Oizumi et al., 2025; Rödström et al., 2022; Su et al., 2012; Ye et al., 2021). To our knowledge, this is the first study of its kind to establish the neuroanatomical basis of these BB biomarkers in rat models of PD, which provides needed biological context for interpreting BB biomarker changes in longitudinal clinical studies and future disease-modifying therapeutic trials.

The increases of the 3 biomarkers in 6-OHDA lesioned rats were above serum levels associated with a sham-operation of the nigrostriatal pathway (Kasanga et al., 2024), strongly suggesting clear that these increases were, at least in part, associated with severe loss of the nigrostriatal pathway and not non-specific tissue injury. Moreover, when CNS levels of these biomarkers in the 6-OHDA cohort were compared against expression levels in the time-matched sham-cohort, the results showed that UCHL-1, GFAP and NfL expression were greater in the SN in the 6-OHDA cohort. In a parallel manner, UCHL-1, GFAP and NfL were also increased in SN in the aged Pink1 KO rats against the aged-matched WT cohort. Like our data, Croucher and colleagues (2025) showed in another genetic rodent PD model (ATP12A2) that GFAP was also increased in the SN as a function of aging (Croucher et al., 2025). Thus, our study gives credence to the idea that blood levels of UCHL-1, GFAP and NfL in human PD should have predictive validity to reflect the severity of nigrostriatal DA neuron loss or loss of DA signaling.

Although changes in GFAP and NfL have been correlative with disease severity in PD (Su et al., 2012; Mollenhauer et al., 2020; Ye et al., 2021; Ygland Rödström et al., 2022; Lin et al., 2023; Tang et al., 2023; Youssef et al., 2023), there is no hard evidence that these biomarker levels corroborate deficiencies in nigrostriatal DA signaling. In humans, any query into this relationship would necessitate the availability of reliable imaging of the same biomarkers in disease-vulnerable CNS tissue to evaluate the level of correlation with blood levels. These quantitative approaches in human are not presently available. This gap underscores a critical unmet need in translational PD research which is the ability to anchor BB biomarker changes to a specific stage of nigrostriatal degeneration. In humans, this will likely require a multimodal approach, such as BB biomarker profiling combined with dopaminergic neuroimaging. Until such approaches are feasible and widely available, well-characterized pre-clinical PD models offer the most tractable system for establishing this biological grounding. As PD is progressive, in both models we captured two different time points to look at biomarker profiles as DA and TH loss occurred at 7 and 28 days for the 6-OHDA model and at ages 7 and 18 months old in the Pink1 KO. In the Pink1 KO, the major risk factor of PD (aging) was also introduced into the neurobiological background. In the 6-OHDA model, the representation of human PD was emulated by the greater rate of TH and DA loss in the striatum compared with the SN in human PD (Kordower et al., 2013). In the Pink1 KO, aging was the critical variable in the study timeline, and although loss of nigrostriatal neurons or TH has been reported inconsistently (Dave et al., 2014; de Haas et al., 2019), we did in fact report an aging-related loss of DA and TH protein in the SN in the samples used for this biomarker study (Soto et al., 2024a).

The SN was the locus of changes observed in expression of the three biomarkers and, critically, common in both models when loss of TH protein was greater at the latter time points. Notably, the level of biomarker expression in the allied tissue of the nigrostriatal pathway was greater in the SN versus the striatum (as seen in the sham-op control group). Given these common differences in biomarker expression pattern between the two PD models, it is plausible that the BB biomarker profile is associated with TH and DA loss in the SN. Indeed, the changes in the serum reflected increases in these biomarkers at the severe stage of nigrostriatal neuron loss in both models for GFAP and NfL, and for UCHL-1 in the Pink1 KO. This is a critical observation in the context of PD, because nigrostriatal pathology begins in the striatum, and the loss of TH and DA is much more severe therein upon disease diagnosis when motor impairments first strike (Bernheimer et al., 1973; Kordower et al., 2013). Primate PD models substantiate that this level of loss in striatumis associated with the onset of motor impairment (Bezard et al., 2001). From a clinical standpoint, this means that by the time a patient presents with motor decline, the window for neuroprotective intervention at the striatal level may be largely closed. The biomarker changes observed here at more moderate (in the Pink1 KO) to severe (in the 6-OHDA model) stages of nigrostriatal loss suggest that GFAP, NfL, and UCHL-1 may be most informative as a tool for monitoring disease progression and therapeutic response after diagnosis.

To this point, despite near complete loss of TH within 5 years of diagnosis (Kordower et al., 2013), the severity of motor impairment continues to worsen (Furukawa et al., 2022; Karimi et al., 2013; Perlmutter and Norris, 2014). Our 6-OHDA model also reflects this clinical observation (Kasanga et al., 2023a). As there still remains anywhere from 10-50% of neurons in the SN at the more advanced stages of the disease (Fearnley and Lees, 1991; Kordower et al., 2013), we speculate that the BB biomarker profile in human PD may reflect expression profiles associated with remaining nigrostriatal neurons in the SN. Thus, if 10–50% of SN neurons remain viable even at moderate-to-advanced disease stages, those surviving neurons represent the biological substrate that a disease-modifying therapy could protect. Moreover, there is substantive preclinical evidence that DA loss in the SN alone is associated with motor decline in aging and in PD models (Kasanga et al., 2023a; Soto et al., 2024a; rev. Salvatore, 2024). Thus, as the clinical results on GFAP and NfL in particular show correlation to disease severity (Su et al., 2012; Mollenhauer et al., 2020; Ye et al., 2021; Ygland Rödström et al., 2022; Lin et al., 2023; Tang et al., 2023; Youssef et al., 2023), we determined if biomarker expression profile in the SN correlated with DA tissue content. UCHL-1 and NfL had positive correlation with DA tissue levels in the SN. GFAP had a negative correlation with remaining TH only in the Pink1 KO model, with greater GFAP expression associated with less TH protein. These data suggest that UCHL-1 and NfL expression are associated with nigrostriatal neuron viability and DA signaling homeostasis in the SN. Thus, the increase in these biomarkers in blood may signify there is loss inviable DA neurons. GFAP expression, on the other hand, appears to increase when DA homeostasis is undermined by loss of neurons or TH protein itself, as would be suggested by the result from the Pink1 KO.

UCHL1, a deubiquitinating (DUB) enzyme, antagonizes ubiquitination of substrates through hydrolase activity, which replenishes the ubiquitin monomer pool for reuse (Snyder, 2021; Bett, 2015; Wilkinson, 1989). Our findings show elevated levels of UCHL1 in aged Pink1 KO rats, suggesting a compensatory response to neuronal damage or elevated levels of protein aggregation. Such an increase would potentiate the efficiency of the ubiquitin-proteasome system in PD pathology (Mi, 2021; Bazarian, 2023; Chen, 2010; Gong, 2006; Zhang, 2014). This possibility is supported by Powis et al (2014) who found pharmacological inhibition of UCHL1 exacerbates disease symptoms in a mouse model of spinal muscular atrophy. We also point out that our results indicate UCHL-1 levels in the SN may be largely driven by nigrostriatal neuron viability. Thus, although total UCHL-1 levels decreased in the SN over time in the 6-OHDA model, it is likely that this decrease was specifically tied to the loss of DA neurons, as UCHL-1 is expressed in nigral DA neurons (Xilouri et al., 2012). As such, when we normalized UCHL-1 to remaining TH protein in both models, there was a dramatic increase at latter stage in the 6-OHDA model (day 28), similar to that found in the aged Pink1 KO. Our results indicate UCHL1 expression increases when TH protein decreases in both models, and that UCHL1 correlates with DA content. Thus, from two PD models, the results point to UCHL1 having a yet unrealized role in DA homeostasis, and its levels may increase in cellular conditions with mitochondrial impairment, as occurs in both models, to preserve nigrostriatal neuron viability and function. Indeed, such a relationship has been recently unveiled in cardiomyocytes (Wu et al., 2022), and in this case, mutations in UCHL-1 exacerbate nigrostriatal neuron loss (Setsuie et al., 2007). Thus, it is plausible that UCHL-1 levels adapts to cellular perturbations affecting mitochondrial function, rather than represent a passive consequence of neuronal damage. Thus, that increased UCHL-1 levels in serum in both models may be indicative of these compensatory changes and it raises the possibility that augmenting this response may confer neuroprotection.

The age-dependent elevation of GFAP in the Pink1 KO, and in the latter stage of neuronal loss in the 6-OHDA model aligns with findings from human studies that show GFAP levels correlate with age, and PD symptom severity (Su et al., 2012; Lin et al., 2023; Youseff et al., 2023). Notably, we previously reported that GFAP levels increase in astrocytes, specifically in the SN, between 7 and 28 days in the 6-OHDA model used herein (Kasanga et al., 2023b). Of particular importance is that this increase was bilateral; thus the increase also occurs in the SN contralateral to lesion. While the precise role of GFAP in the pathology of PD remains to be fully elucidated, research involving GFAP-deficient mice demonstrated an increase in glial-derived neurotrophic factor (GDNF), a neuroprotective agent that regulates DA signaling and has therapeutic potential for PD (Hanbury et al., 2003; Brenner, 2014; Kiri et al., 2001; Mätlik et al., 2018; Pertusa et al., 2008; Manfredsson et al., 2020; Sandoval et al., 2026). The results from this study, alongside the established correlation between serum GFAP levels and motor/cognitive performance in PD patients, underscore GFAP’s utility as a multifaceted marker that is a critical component of a biomarker profile to delineate PD progression, as increased levels in the SN and serum may correspond to the severity of TH protein and DA loss in the SN.

NfL has long been known as an indicator of brain injury and neurodegeneration. Its relevance as a biomarker comes from results showing its levels in blood correspond to axonal damage (Bäckström et al., 2020; Mollenhauer et al., 2020). In the 6-OHDA and Pink1 KO models NfL increased with decreasing TH protein levels in the SN, suggesting that elevated amounts of NfL in serum resulted from loss of cell membrane from the nigrostriatal neurons. The release of NfL from damaged neurons is thought to be facilitated by calpain, a calcium-dependent cysteine protease implicated both in neuronal remodeling and axon degeneration (Kahn et al., 2025; Metwally et al., 2021). Initial insults to DA neurons in PD have been theorized to come from dysregulation of neuronal calcium homeostasis and excess exposure to glutamate, leading to excitotoxicity (Chotibut et al., 2014; Kandy et al., 2022; Wu et al., 2025). There is evidence for excitotoxicity in PD models, specifically in both striatum and the SN, and increasing glutamate uptake can mitigate loss of TH (Chotibut et al., 2014). Excessive levels of fragmented NfL can be produced by damaged neurons in a simulated excitotoxic environment, triggering release levels above a normal baseline. This, in turn, can provoke a pro-inflammatory immune response from nearby microglia, exacerbating neuronal injury and cell death (Kahn et al., 2025). The immune-activating nature of NfL marks it as both a potential biomarker, and a potential therapeutic target. In fact, the FDA recently approved a new drug for amyotrophic lateral sclerosis, which effectively lowers plasma NfL levels (Benatar, 2022; 2023). Additionally, inhibition of NfL uptake by microglia via anti-NfL antibodies can reduce the pro-inflammatory microglia response, suggesting that targeting NfL could prevent immune-mediated inflammation in neurodegenerative disorders, like PD and ALS (Kahn et al., 2025). Overall, our findings support that NfL is a critical component of the biomarker profile that can be used to gauge severity of dysfunction in nigrostriatal DA signaling, particularly in that NfL levels rise against TH protein loss in the SN, and are associated with increased serum levels, in both models.

### Conclusion

Our results have addressed a critical knowledge gap in our understanding of whether changes in blood levels could reflect nigrostriatal DA signaling. Using two different PD models at two different time points after lesion induction or age, our results point to the possibility that changes in BB biomarkers UCHL-1, GFAP, and NfL, together may be indicative of the functional capacity of DA signaling (Kasanga et al., 2023, 2024; Soto et al., 2024a; Galfano et al., 2026). Critically, these changes in expression in both the CNS and serum aligns with the severity of TH protein loss only in the SN. Accordingly, our study makes a significant advance to provide some measure of face validity that serum biomarker levels may gauge nigrostriatal neuronal loss and impaired DA signaling, as previously presumed in clinical studies showing correlation between serum levels of NfL and GFAP against PD symptom severity. Moreover, the alignment of biomarker changes in the SN may have functional relevance, given the clinical correlations, in that nigral DA signaling alone can affect locomotor function in preclinical studies (Emborg et al., 1998; Salvatore, 2024 (rev)) and is supported by clinical observations (Kordower et al., 2013; Perlmutter and Norris, 2014; Furukawa et al., 2022). It is yet unknown how UCHL-1, GFAP, and NfL levels may change in other brain regions affected in PD pathology, such as the locus ceruleus and prefrontal cortex, pathologies within which correspond to deficits in executive function at the early stages of the disease (Nejtek et al., 2021; Salvatore et al., 2021; Doshier et al., 2025), including in the Pink1 KO model used herein (Soto et al., 2024b). One final highlight is that there is consistency between the two rodent models and human PD, in that TH loss in the SN is more protracted, compared to striatum; giving credence to the idea that our findings may extend to human PD. To this point, we found each biomarker was expressed at greater levels in the SN versus striatum, suggesting a greater relative contribution to the blood from SN. Our study is an advance toward face validity that the levels of the BB biomarkers UCHL-1, GFAP, and NfL can gauge severity of the loss of nigrostriatal DA signaling, in addition to correlation with motor impairment as previously shown in clinical studies. It provides biological grounding for the use of UCHL-1, GFAP, and NfL as a composite biomarker panel that may reflect nigrostriatal integrity by tracking the biological status of surviving nigral neurons rather than simply marking cumulative loss. Additionally, our results also indicate that compensatory responses, such as UCHL-1 upregulation, may slow nigrostriatal degeneration, and represent an important avenue for future investigation

## Supporting information

Supplemental Results

## Acknowledgements

This work was supported by the Department of Defense Parkinson’s Research Program, Investigator-Initiated Research Award (W81XWH-19-1-0757) award to MFS and R01ES033892 and RF1NS130713 to JRR. IS was supported by the National Institutes of Health/National Institute on Aging Predoctoral International Fellowship (T32 AG020494).

## Notes

### Competing Interest Statement

The authors have declared no competing interest.

