## Supplemental Results for "Serum UCHL-1, GFAP, and NfL track tyrosine hydroxylase loss in substantia nigra in two Rat Models of Parkinson’s Disease"

**Supplemental Figures**

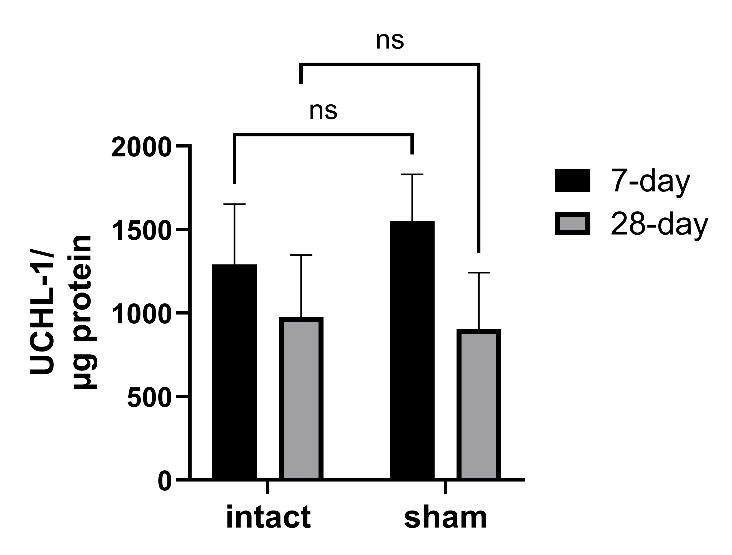

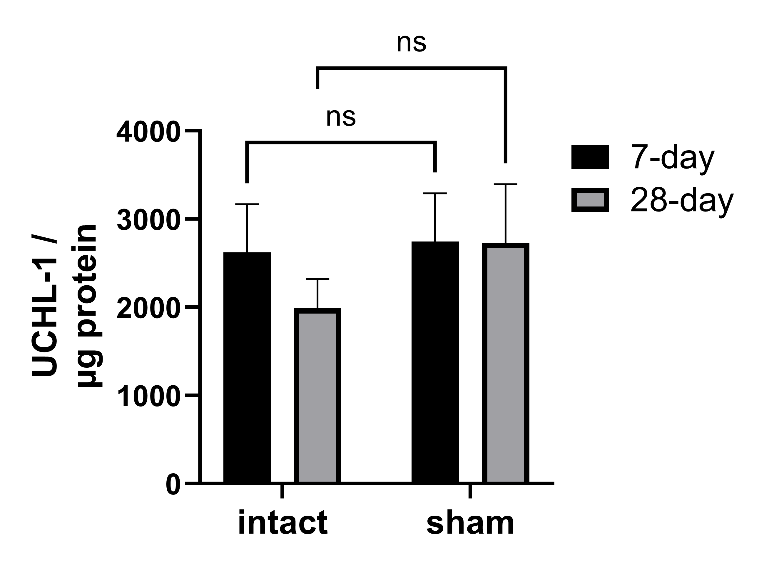
 **SN	striatum**

**Figure S1. UCHL-1 expression after sham-operation. SN.** Sham-operation did not affect UCHL-1 expression at either time point after surgery. Sham-op (F(1,10)=0.76, *p=*0.40), days past sham-op (F(1,13)=0.28, *p=*0.61), interaction (F(1,10)=0.29, *p=*0.60). **striatum.** Sham-operation did not affect UCHL-1 expression at either time point after surgery. Sham-op (F(1,12)=0.03, *p=*0.86), days past sham-op (F(1,13)=0.84, *p=*0.37), interaction (F(1,12)=0.42, *p=*0.53).

**SN	striatum**

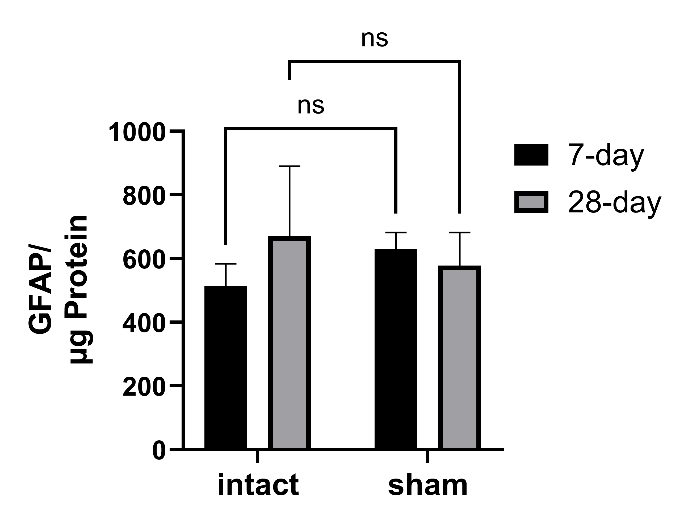

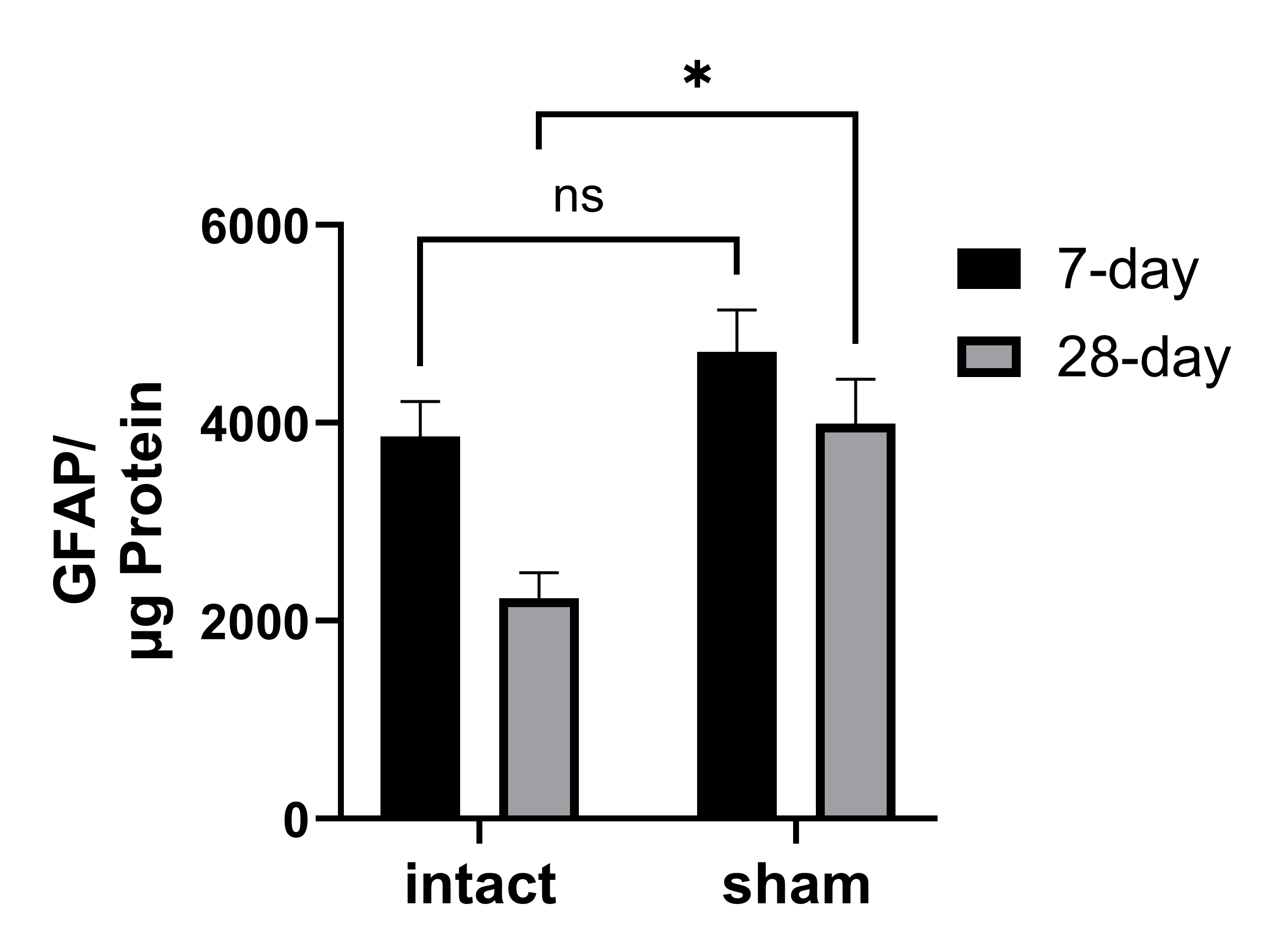

**Figure S2. GFAP expression after sham-operation. SN.** Sham-operation affected GFAP expression, with lower expression levels in the intact side at day 28. Sham-op (F(1,8)=11.9, *p=*0.009), days past sham-op (F(1,13)=9.33, *p=*0.009), interaction (F(1,8)=1.44, *p=*0.26). Day 28, intact v sham (t=3.10, **p*=0.03). **striatum.** Sham-operation did not affect GFAP expression at either time point after surgery. Sham-op (F(1,10)=0.03, *p=*0.87), days past sham-op (F(1,11)=0.11, *p=*0.74), interaction (F(1,10)=2.18, *p=*0.17). Between the SN and striatum, GFAP expression levels were ~5- to 10-fold greater in the SN.

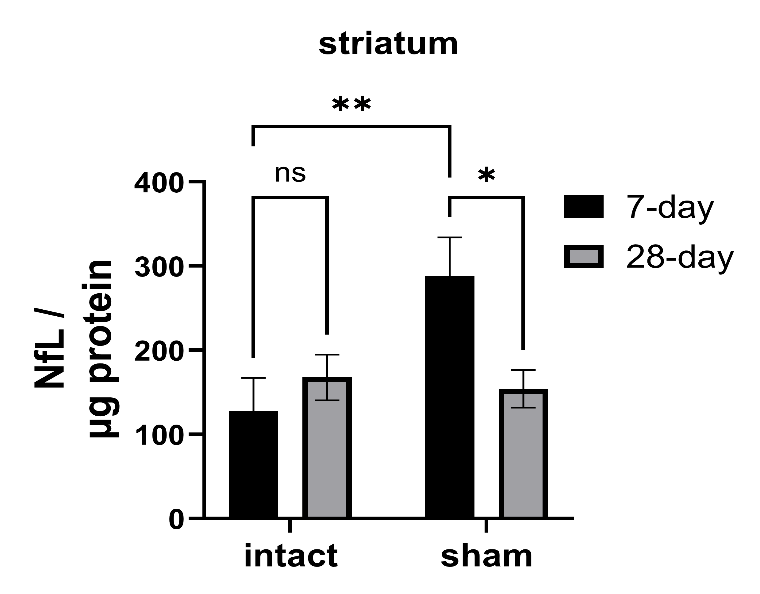

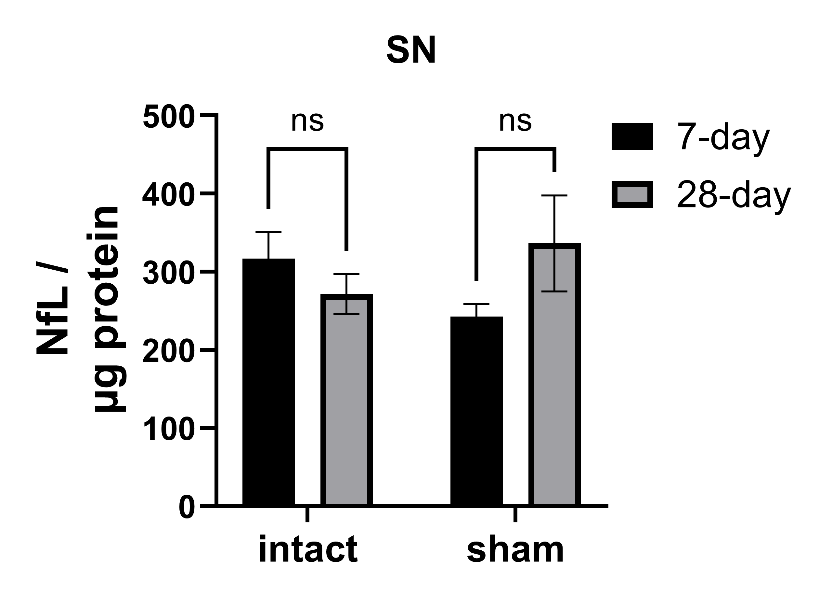
**Suppl Fig 3. NfL expression.**

**Figure S3. NfL expression after sham-operation. SN.** Sham-operation had no effect on NfL expression in SN. Sham-op (F(1,9)=0.02, *p=*0.89), days past sham-op (F(1,12)=0.26, *p=*0.62), interaction (F(1,9)=3.6, *p=*0.09). **Striatum.** Sham-operation transiently increased NfL expression early after surgery (F(1,21)=4.46, *p=*0.047), days past sham-op (F(1,21)=1.81, *p=*0.19), interaction (F(1,21)=6.26, *p=*0.021). Sham; *day 7 v day 28 (t=2.67, *p*=0.01); 7 day, intact v sham (t=3.20, ***p*=0.004).
